# Transcriptional reprogramming deploys a compartmentalized “timebomb” in *Catharanthus roseus* to fend off chewing herbivores

**DOI:** 10.1101/2024.06.11.598499

**Authors:** Yongliang Liu, Jizhe Shi, Barunava Patra, Sanjay Kumar Singh, Xiaoyu Liu, Yongqing Li, Ying Wang, Xuguo Zhou, Sitakanta Pattanaik, Ling Yuan

**Affiliations:** Department of Plant and Soil Sciences and Kentucky Tobacco Research and Development Center, University of Kentucky, Lexington, Kentucky 40546; Department of Entomology, University of Kentucky, Martin-Gatton College of Agriculture, Food and Environment, Lexington, Kentucky 40546; Pomology Institute, Shanxi Agricultural University, Taigu 030815, Shanxi, China; Guangdong Provincial Key Laboratory of Applied Botany, South China Botanical Garden, Chinese Academy of Sciences, Guangzhou 510520, China; Department of Entomology, School of Integrative Biology, College of Liberal Arts & Sciences, University of Illinois Urbana-Champaign, Urbana, IL 61801, USA

**Keywords:** *Catharanthus roseus*, terpenoid indole alkaloid, insect-plant interaction, transcriptional reprogramming, timebomb, herbivory resistance, cold tolerance

## Abstract

The evolutionary arms-race between plants and insects has led to key adaptive innovations that drive diversification. Alkaloids are well-documented anti-herbivory compounds in plant chemical defenses, but how these specialized metabolites are allocated to cope with both biotic and abiotic stresses concomitantly is largely unknown. To examine how plants prioritize their metabolic resources responding to herbivory and cold, we integrated dietary toxicity bioassay in insects with co-expression analysis, hierarchical clustering, promoter assay, and protein-protein interaction in plants. *Catharanthus roseus,* a medicinal plant known for its insecticidal property against chewing herbivores, produces two terpenoid indole alkaloid monomers, vindoline and catharanthine. Individually, they exhibited negligible toxicity against *Manduca sexta,* a chewing herbivore; their condensed product, anhydrovinblastine, however, was highly toxic. Such unique insecticidal mode of action demonstrates that terpenoid indole alkaloid “timebomb” can only be activated when the two spatially isolated monomeric precursors are dimerized by herbivory. Without initial selection pressure and apparent fitness costs, this adaptive chemical defense against herbivory is innovative and sustainable.

The biosynthesis of insecticidal terpenoid indole alkaloids is induced by herbivory but suppressed by cold. Here, we identified a transcription factor, <u>H</u>erbivore Induced <u>V</u>indoline-gene <u>E</u>xpression (HIVE), that coordinates the production of terpenoid indole alkaloids in response to herbivory and cold stress. The HIVE-mediated transcriptional reprogramming allows this herbaceous perennial to allocate their metabolic resources for chemical defense at a normal temperature, when herbivory pressure is high, but switches to cold tolerance under a cooler temperature, when insect infestation is secondary.

## Introduction

The arms race between plants and their insect herbivores has co-evolved for more than 400 million years (Kareiva, 1999; Price, Denno, Eubanks, Finke, & Kaplan, 2011), at morphological (Atsatt & O’Dowd, 1976; Hanley, Lamont, Fairbanks, & Rafferty, 2007), molecular (Atsatt & O’Dowd, 1976; Erb & Reymond, 2019; Hanley et al., 2007; Xia et al., 2021), and biochemical levels (Fürstenberg-Hägg, Zagrobelny, & Bak, 2013; Mithöfer & Boland, 2012). During the interaction with insect herbivores, plants have evolved strategies that directly (e.g., apparency, structural, and chemical) and indirectly (e.g., “calling for help”-plants emit herbivore-induced volatiles to recruit predators when attacked) defend against herbivory. Plant specialized metabolites are the most effective and diverse insecticidal chemicals involved in defense through either qualitative or quantitative means. Most qualitative defenses occur in tissues that are vulnerable over short time spans, such as young, tender leaves or seeds, and are effective on most generalists and often recycled when no longer needed.

Specialized metabolites are essential for the communication between plants and their surrounding environments, especially under biotic and abiotic stress conditions. Specialized metabolites represent a rich reservoir for novel compounds with pharmaceutical/therapeutic and pest/disease management implications. The medicinal plant *Catharanthus roseus* produces almost 200 terpenoid indole alkaloids (TIAs), and among them, the anticancer drugs vinblastine and vincristine are widely used in various chemotherapies (Roepke et al., 2010). Strictosidine serves as a universal precursor for various TIAs, including ajmalicine, catharanthine, tabersonine, and vindoline (Figure S1a). Furthermore, catharanthine and vindoline are condensed to produce the dimeric TIAs vincristine and vinblastine, which are highly toxic to insects. The precursors (catharanthine and vindoline) were initially suggested to accumulate strictly in distinct cellular compartments of *C. roseus* leaves. Upon cell damages caused by insect attack, catharanthine and vindoline converge, facilitating enzymatic condensation to form the dimeric vincristine and vinblastine (Roepke et al., 2010). Recent single cell metabolome analyses have indicated that catharanthine and vindoline co-localize in idioblast and laticifer cells (Guedes et al., 2024; Li et al., 2023; Yamamoto et al., 2019; Yamamoto et al., 2016). Despite their co-localization, the concentrations of the dimeric compounds remain low, raising another possibility that the enzymes responsible for dimerization are localized in cell types other than idioblast and laticifer. One evidence supporting this possibility is that CrPRX1, a peroxidase involved in dimerization of catharanthine and vindoline, is selectively expressed in the epidermis (Costa et al., 2008). Both concepts, either compartmentalization of precursors or separation of enzyme and substrates, suggest a compartmentalized “Timebomb” against herbivores, similar to the classic “mustard oil bomb”, in which the precursors and enzymes related to the synthesis of primary defensive chemicals in plants are spatially separated (Lüthy & Matile, 1984; Matile, 1980). Documented herbivory by insects on *C. roseus* is rare and is mainly limited to insects with piercing-sucking mouthparts, whereas dietary toxicity has been reported in a variety of chewing insect herbivores (Dugé de Bernonville et al., 2017; Meisner, Weissenberg, Palevitch, & Aharonson, 1981; Roepke et al., 2010). The mode of action of such a “timebomb” against chewing insects remains largely unknown.

In addition to adaptations to biotic stresses, plants have evolved sophisticated mechanisms to cope with abiotic stresses, including cold, drought, and salt. Cold stress is one of the key environmental factors that impairs the growth and productivity of plants (Sanghera, Wani, Hussain, & Singh, 2011). TIA contents in *C. roseus* leaves drop significantly following exposure to cold treatment (Dutta, Sen, & Deswal, 2007). Cold stress also negatively affects insect activity (Mellanby, 1939). As the threat of herbivory declines, nitrogen stored in plant metabolites is recycled for use in plant growth/cold acclimation (Poulton, 1990). The underlying mechanisms of how plants balance their metabolic efforts on cold tolerance and herbivory defenses remain poorly understood. In this study, we focused on how the plant balances the production of catharanthine and vindoline to manage biotic (herbivory) and abiotic (cold) stresses. Building on previous knowledge and our preliminary research, we hypothesized that *C. roseus* fine-tunes its stress responses, specifically herbivory resistance and cold tolerance, through transcriptional reprogramming of plant specialized metabolism. To examine this overarching hypothesis, we addressed the following questions: 1) mode of action of the TIA “timebomb”, a qualitative chemical defense in *C. roseus* against chewing herbivores, 2) crosstalk between chemical defense and cold tolerance in *C. roseus*, and 3) transcriptional reprogramming of TIA biosynthesis in response to herbivory- and cold-induced stresses.

## Results

### Mode of action of the compartmentalized catharanthine-vindoline “timebomb”

#### Toxicity

When treated with vindoline, catharanthine, and mixtures of the two at different concentrations (0.0219, 0.219, and 2.19 μmol/ml, respectively; Figure 1c), larval mortality of tobacco hornworm (*Manduca sexta*) was significantly different between the three alkaloid treatments at 24, 48, 72, and 96 h (Figure 1a). The mixture showed higher dietary toxicity on *M. sexta* larvae than did either of the two individual compounds. Acute toxicity was only found in the mixture rather than in the two individual compounds. Catharanthine and vindoline showed synergistic toxicity in *M. sexta* larvae across all concentrations at all time points (Table 1; *χ*^2^ > 3.84 with *df* = 1 and *p* = 0.05).

**Figure 1:**
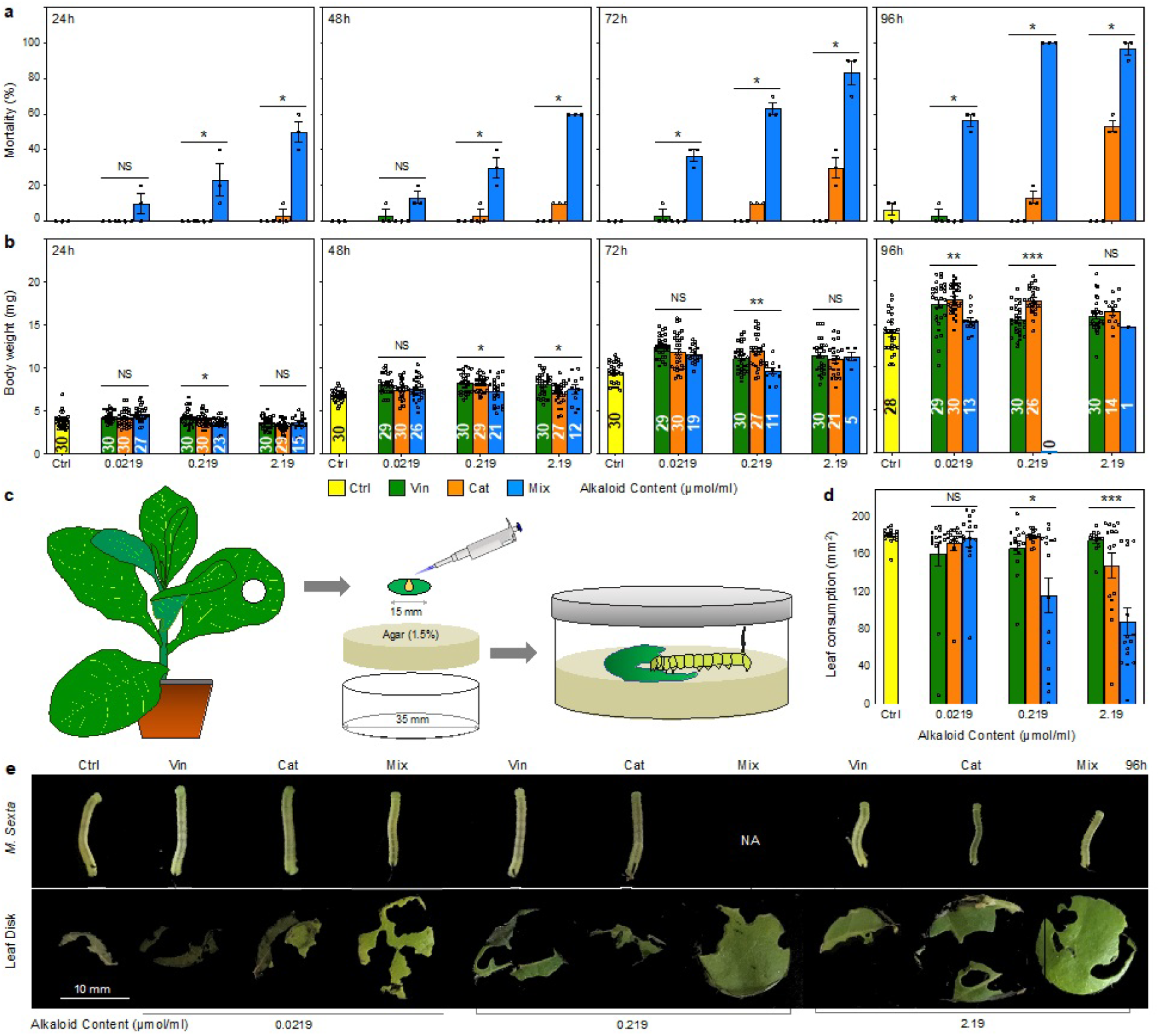
Dietary effects of the two TIAs and the mixture on *M. sexta* larvae. **a** and **b** Mortality and body weights in the different treatments at three TIA concentrations at 24, 48, 72, and 96 h after larval transfer. **c** Design of the *M. sexta* feeding bioassay using TIA-treated tobacco leaf disks. **d** Leaf consumption in the different treatments at the three TIA concentrations at 24, 48, 72, and 96 h. **e** *M. sexta* larvae after 96 h of feeding on TIA-supplemented leaf disks (upper) and the remaining tobacco leaf disks (lower). Scale bar = 10 mm. In **a**, **b**, and **d**, the different colors represent the three TIA treatments. The small circles indicate the data distribution within each treatment. Data are presented as mean ± SE (in **a**, n = 3; in **b**, n is labeled in each bar; in **d**, n=15). NS represents no significant difference, \**p* <0.05, \*\**p* <0.01, \*\*\**p* <0.001; in **a** and **d**, the Kruskal-Wallis test was used, while in **a** one-way ANOVA was used for comparisons.

**Table 1.** Synergistic effect of terpenoid indole alkaloid binary mixture against *Manduca sexta*.

| Concentration<br>(nmol/mL) | Assay<br>Time (h) | Mortality (%) | | | | $\chi^2$ | Ratio* | Effect** |
| --- | --- | --- | --- | --- | --- | --- | --- | --- |
|  |  | Pure Compound |  | Binary Mixture |  |  |  |  |
|  |  | Catharanthine | Vindoline | Expected | Observed |  |  |  |
| 2.19 | 24 | 3.33 | 0.00 | 3.33 | 50.00 | 653.33 | 15.00 | Synergistic |
| 2.19 | 48 | 10.00 | 0.00 | 10.00 | 60.00 | 250.00 | 6.00 | Synergistic |
| 2.19 | 72 | 30.00 | 0.00 | 30.00 | 83.33 | 94.81 | 2.78 | Synergistic |
| 2.19 | 96 | 53.33 | 0.00 | 53.33 | 96.67 | 35.21 | 1.81 | Synergistic |
| 0.219 | 24 | 0.00 | 0.00 | 0.00 | 23.33 | NA | NA | Synergistic |
| 0.219 | 48 | 3.33 | 0.00 | 3.33 | 30.00 | 213.33 | 9.00 | Synergistic |
| 0.219 | 72 | 10.00 | 0.00 | 10.00 | 63.33 | 284.44 | 6.33 | Synergistic |
| 0.219 | 96 | 13.33 | 0.00 | 13.33 | 100.00 | 563.33 | 7.50 | Synergistic |
| 0.0219 | 24 | 0.00 | 0.00 | 0.00 | 10.00 | NA | NA | Synergistic |
| 0.0219 | 48 | 0.00 | 3.33 | 3.33 | 13.33 | 30.00 | 4.00 | Synergistic |
| 0.0219 | 72 | 0.00 | 3.33 | 3.33 | 36.67 | 333.33 | 11.00 | Synergistic |
| 0.0219 | 96 | 0.00 | 3.33 | 3.33 | 56.67 | 853.33 | 17.00 | Synergistic |

\*Ratio = Observed/Expected.
\*\* $\chi^2 > 3.84$ denotes synergistic effect.

#### Body weight

Body weights of *M. sexta* larvae were influenced by dietary treatments containing catharanthine, vindoline, and the mixture of the two at three different concentrations for 96 h (Figure 1b). At 24 h, as the concentrations increased, body weights decreased slightly in the different treatments, with no difference from the control (except when treated with the mixture at 0.0219 μmol/ml). There was a significant interaction between treatment and concentration (*F*(6, 333) = 3.92, *p* <0.001). At 48 h, larval body weight was mainly influenced by the treatment (*F*(3, 323) = 22.25, *p* <0.001) with no interaction between treatment and concentration (*F*(6, 323) = 1.79, *p* = 0.10). When treated with vindoline at all concentrations and catharanthine at concentration of 0.219 μmol/ml, body weight was significantly higher than in the control, while there were no differences between all the mixture treatments and the control (*p* >0.05). At 72 h, both treatment (*F*(3, 291) = 36.51, *p* <0.001) and concentration (*F*(2, 291) = 6.52, *p* <0.01) influenced body weight, with an interaction effect (*F*(6, 291) = 2.94, *p* < .01). Except for the treatments with the mixture at concentrations of 0.219 and 2.19 and catharanthine at 2.19 μmol/ml, body weight was higher than in the control in all other treatments across the different concentrations (Tukey-Kramer HSD test, *p* <0.001). At 96 h, the body weight was influenced by treatment (*F*(2, 256) = 57.40, *p* 0.001), with no interaction with concentration (*F*(5, 256) = 2.01, *p* = 0.08). The body weights of larvae treated with the mixture at concentrations of 0.0219 and 2.19, or vindoline at 0.219 μmol/ml, showed no differences compared to the control (Tukey-Kramer HSD test, *p* >0.05). After treatment with the vindoline/catharanthine mixture at 2.19 μmol/ml for 96 h, only one larva survived, while no survivor at the 0.219 μmol/ml treatment. The sole survivor could be an outlier.

#### Leaf consumption

Leaf consumptions were significantly different across the different treatments (Figure 1d and 1e). When treated with the vindoline/catharanthine mixture at 0.219 and 2.19 μmol/ml, larvae consumed significantly less than those in the other treatments (Wilcoxon signed-rank test; *p* <0.05), with no differences between the treatment concentrations (Z = 1.64, *p* = 0.10). There was no difference in the starting areas of the leaf disks used in all the treatments (ANOVA; *F*(9, 149) = 1.71, *p* = 0.09).

### Crosstalk between chemical defense and cold tolerance in *C. roseus*

#### The effect of cold temperature on vindoline biosynthesis

Accumulations of catharanthine and vindoline were significantly reduced at low temperature (LT; 4°C) compared to normal temperature (NT; ∼25°C) (Figure 2a; Figure S1b). Tabersonine content was not significantly altered by LT. Expression of five genes (*T16H2*, *T3O*, *NMT*, *D4H*, and *DAT*) associated with vindoline pathway decreased after 2h and 6h of LT treatment in mature leaves and seedlings of *C. roseus*, respectively, compared to the 0-hr control (Figure 2b; Figure S1c).

**Figure 2:**
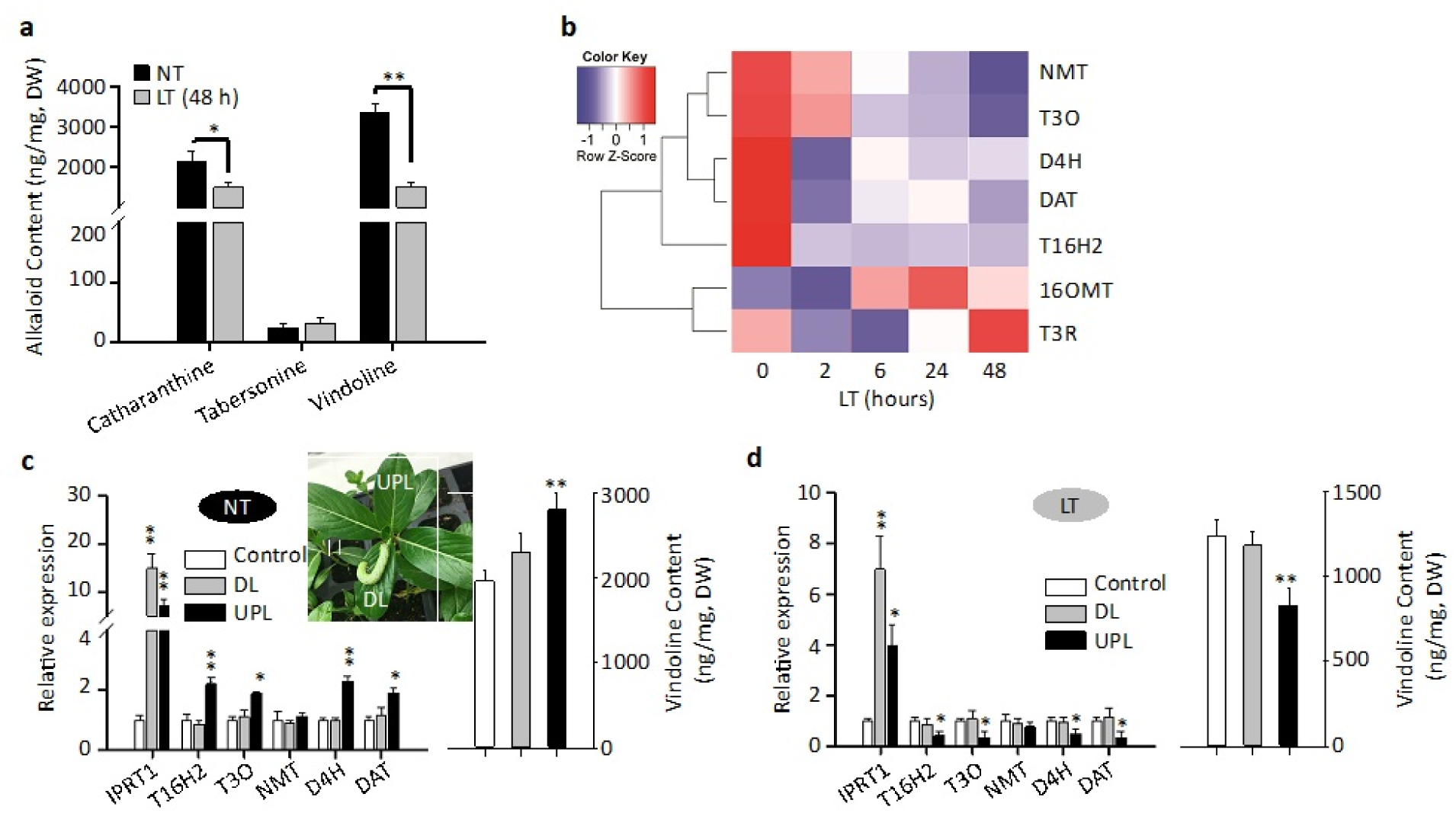
Vindoline biosynthesis is suppressed by low temperature and induced by herbivory. **a** Measurements of catharanthine, tabersonine, and vindoline in *C. roseus* leaves. *C. roseus* plants grown at normal temperature (NT) were either maintained at NT (control) or exposed to low temperature (LT; 4℃) for 48 hours. Alkaloids were extracted and analyzed by LC-MS/MS, and the concentration of each alkaloid was estimated based on peak areas compared with standards. DW, dry weight. **b** Relative expression of seven vindoline pathway genes (*T16H2*, *16OMT*, *T3O*, *T3R*, *NMT*, *D4H*, and *DAT*) in *C. roseus* leaves exposed to LT for different lengths of time (2, 6, 24, 48 hours). Relative gene expression was measured by RT-qPCR and visualized using a heatmap generated with the R package. **c** Vindoline content and relative expression of the herbivory-induced marker gene, *IPRT1,* and five cold-regulated vindoline pathway genes in *C. roseus* leaves two days after *Manduca sexta* herbivory at NT. In the picture, the leaf on which the *M. sexta* larvae is feeding is the damaged leaf (DL), while the paired leaf of the DL is called the undamaged paired leaf (UPL). **d** Vindoline content and relative expression of *IPRT1* and five cold-regulated vindoline pathway genes in leaves (control, DL, and UPL) of *C. roseus* plants exposed to LT for 48 hours after *M. sexta* leaf feeding. Gene expression was measured by RT-qPCR. The values represent means ± SD from three biological replicates. Statistical significance was calculated using Student’s *t* test (* *p* <0.05 and ** *p* <0.01).

#### The effect of herbivory on vindoline biosynthesis

*Intracellular pathogenesis-related protein T1* (IPRT1) gene expression is induced by herbivory in *C. roseus* leaves (Dugé de Bernonville et al., 2017). *IPRT1* expression was significantly induced in the locally damaged leaves (DL) and the undamaged paired leaves (UPL) after herbivory at both NT and LT (Figure 2c, d). Vindoline content, as well as expression of four vindoline pathway genes (*T16H2*, *T3O*, *D4H,* and *DAT*) increased in UPL but not in DL at NT (Figure 2c). However, at LT, vindoline content and the expression of *T16H2*, *T3O*, *D4H,* and *DAT* were significantly decreased in the UPL, but not in the DL, compared to the control (Figure 2d).

### Transcriptional reprogramming of TIA biosynthesis in response to herbivory- and cold-stresses

#### Herbivore Induced Vindoline-gene Expression (HIVE), an activator of vindoline biosynthesis

We focused on the bHLH TF family in *C. roseus* because (1) the E-box motifs, binding sites for bHLH TFs, are present in four (*T16H2*, *T3O*, *D4H*, and *DAT*) of the five cold-responsive vindoline pathway gene promoters, and (2) bHLH TFs are known regulators of plant specialized metabolism and cold tolerance (Gao & Dubos, 2023; Hao, Zong, Ren, Qian, & Fu, 2021). We identified a total of 108 putative bHLH TF genes in the *C. roseus* genome. Thirteen of which were found to be tightly co-expressed with genes associated with the vindoline pathway (Figure S2a; Table S1). Of the thirteen co-expressed bHLH genes, only one, *CRO_T110248*, belongs to sub-group IIIb bHLH (Figure S2b). The subgroup IIIb bHLHs contain regulators of specialized metabolism in *Artemisia annua* and *Phalaenopsis* (Chuang, Hung, Tsai, Chen, & Chen, 2018; Xiang et al., 2019; Yu, Zhang, da Silva, Zhao, & Duan, 2021) and cold tolerance. Transient overexpression of *CRO_T110248* (*CRO_T110248*-OX) resulted in a 2 to 3-fold increase in the expression of *T16H2*, *T3O*, *D4H*, and *DAT* compared to the empty-vector control (EV) (Figure 3a). The vindoline content increased significantly in *CRO_T110248*-OX seedlings compared to that in the EV control (Figure 3a). When *CRO_T110248* expression was down-regulated by VIGS, transcript levels of *T16H2*, *T3O*, *D4H*, and *DAT* were reduced by 25%-70% compared to EV (Figure 3b). Vindoline content was reduced by 37% in leaves of the *CRO_T110248*-VIGS plants compared to EV (Figure 3b). Seedlings of *C. roseus,* in which *CRO_T110248* was co-expressed with individual vindoline gene promoter-*GUS* constructs, showed significantly increased GUS activities compared to seedlings expressing the promoter-*GUS* constructs alone (Figure 3c). The *D4H* promoter (pro*D4H,* 704 bp) contains two bHLH TF binding motifs (E-boxes, 5’-CANNTG-3’; Figure 3d). Mutations in the individual E-boxes or both had no effect on CRO_T110248-mediated activation of pro*D4H* (Figure 3d), indicating that CRO_T110248 indirectly regulates *D4H*.

**Figure 3:**
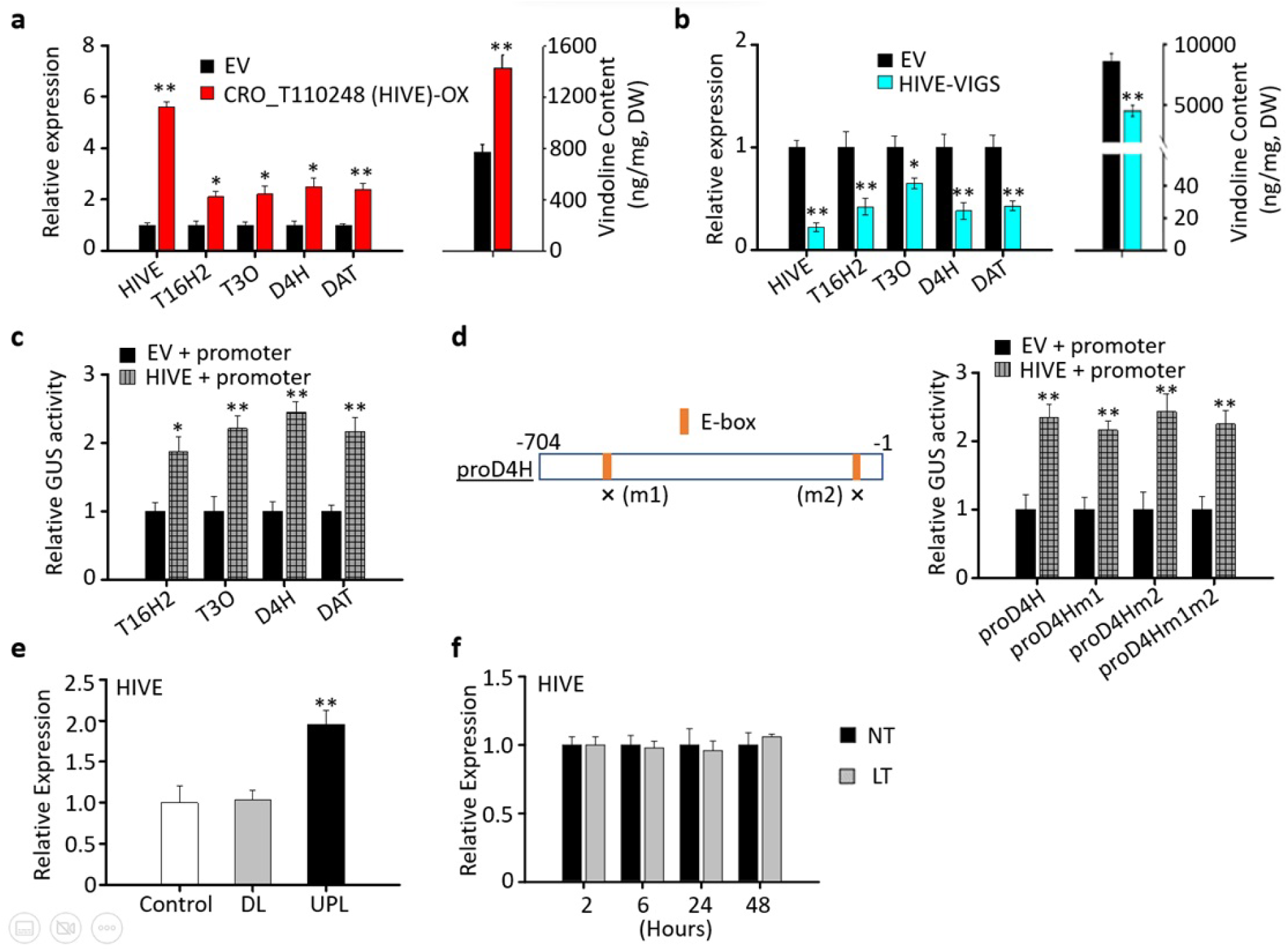
HIVE activates vindoline biosynthesis. **a** Vindoline content and the relative expression of *CRO_T110248* (*HIVE*), *T16H2*, *T3O*, *D4H,* and *DAT* in the empty vector (EV) control and *HIVE*-overexpressing (*HIVE*-OX) *C. roseus* seedlings. **b** Vindoline content and relative expression of *HIVE*, *T16H2*, *T3O*, *D4H,* and *DAT* in the EV and *HIVE*-VIGS leaves of *C. roseus* seedlings. **c** Transactivation of the *T16H2*, *T3O*, *D4H*, and *DAT* promoters by HIVE in *C. roseus* seedlings. **d** Mutation of the E-box motifs has no effect on activation of the *D4H* promoter by HIVE. Schematic diagram on the left showing the E-box motifs in the *D4H* promoter (pro*D4H*). The E-box motifs CACATG (E-box 1) and CAATTG (E-box 2) were mutated (m1 and m2, respectively) to AAAAAA. The effector and reporter (promoter-*GUS*) constructs were co-infiltrated into *C. roseus* seedlings. A plasmid containing the firefly *luciferase* (*LUC*) reporter was used as an internal control. GUS activity was normalized against luciferase activity. The control is the reporter with EV. **e** Relative expression of *HIVE* in *C. roseus* leaves (control, DL, and UPL, as described in Figure 2d) after *M. sexta* herbivory. **f** Relative expression of *HIVE* in *C. roseus* seedlings exposed to LT (4℃) for 2, 6, 24, and 48 hours. Gene expression was measured by RT-qPCR. Alkaloids were extracted and analyzed by LC-MS/MS, and the concentrations of the alkaloids were estimated based on peak areas compared with standards. DW, dry weight. The data represent means ± SD from three biological replicates. Statistical significance was calculated using Student’s *t* test (* *p* <0.05 and ** *p* <0.01).

In addition, we measured the expression of *CRO_T110248* in *C. roseus* leaves after herbivory. *CRO_T110248* expression was not affected in DL but was induced in UPL after herbivory by *M. sexta* (Figure 3e). In addition, the expression of *CRO_T110248* remained unaffected by LT (Figure 3f). CRO_T110248 hereafter designated as Herbivore Induced Vindoline-gene Expression (HIVE).

#### HIVE regulates vindoline biosynthesis through CrMYBR2

To identify critical *cis*-elements in the *D4H* promoter for the activation by HIVE, we generated a truncated *D4H* promoter (pro*D4Hs*; Figure 4a) which excludes the two E-boxes. Like pro*D4H* (Figure 3d), pro*D4Hs* was activated by HIVE in *C. roseus* seedling (Figure 4a). Five different TF binding motifs were identified in pro*D4Hs* (Table S2), including a MYB-related (MYBR) TF binding site, the CR-box (CCA1-like/R-R-type binding box). In addition to a CR-box motif in pro*D4Hs*, CR-boxes were also identified in the promoters of *T16H2* (GATAAA, GATATT), *T3O* (GATATA), and *DAT* (GATAAT). The activation of pro*D4Hs* or pro*D4H* by HIVE decreased significantly when the CR-box sequence was mutated (pro*D4Hsm3* or pro*D4Hm3*) (Figure 4b), suggesting HIVE regulates the promoter activity indirectly through one or more CCA1-like/R-R-type MYBR factors.

**Figure 4:**
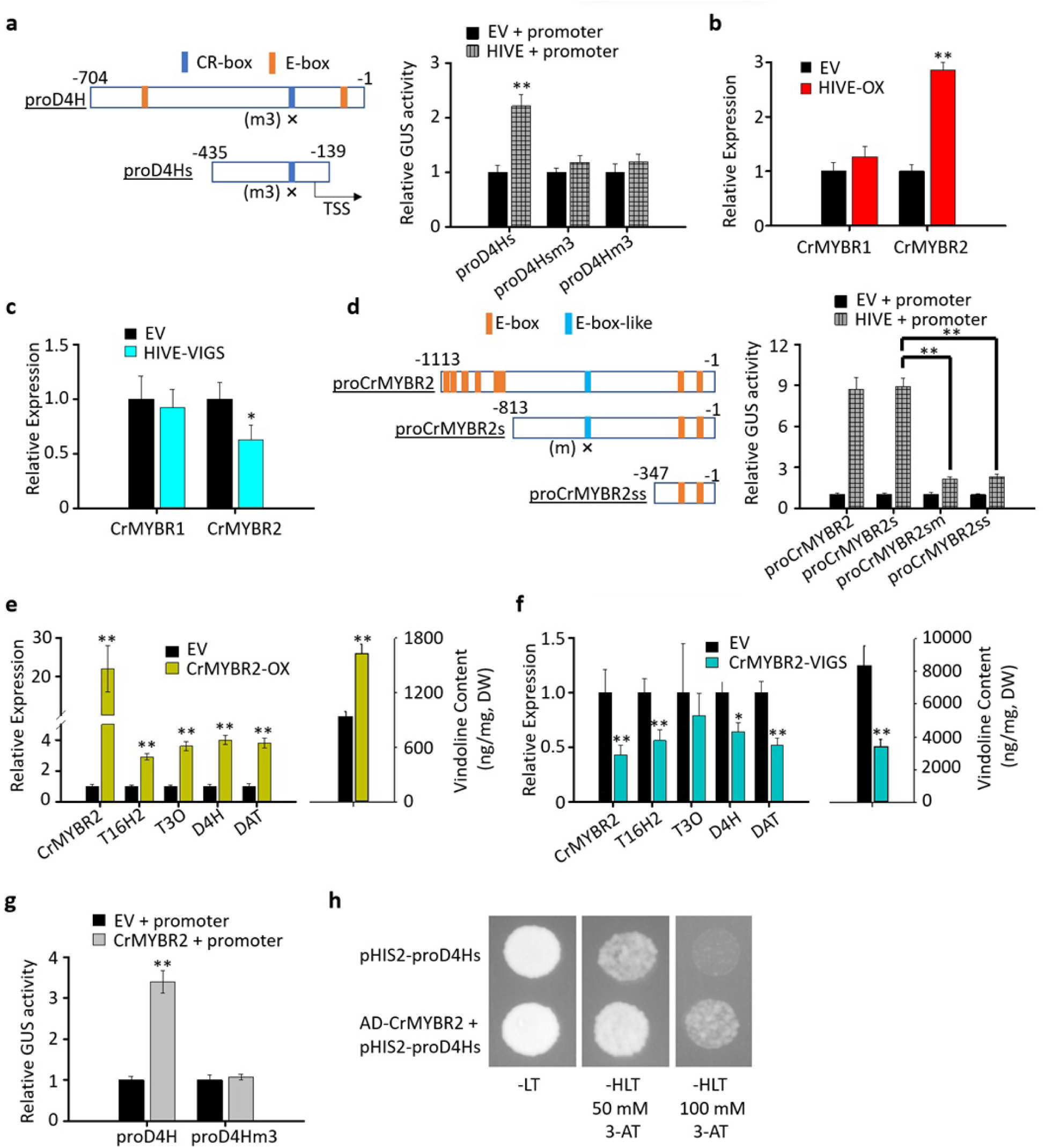
HIVE regulates vindoline biosynthesis through CrMYBR2. **a** Mutation of the CR-box motif negatively affects activation of the full-length (pro*D4Hm3*) and truncated (pro*D4Hsm3*) *D4H* promoter by HIVE. Schematic diagram on the left showing the CR-box motif in the *D4H* promoter (pro*D4H*). The CR-box motif GATATT was mutated to AAAAAA (m3). **b** Relative expression of *CrMYBR1* and *CrMYBR2* in EV and *HIVE*-OX *C. roseus* seedlings. **c** Relative expression of *CrMYBR1* and *CrMYBR2* in EV and *HIVE*-VIGS leaves. **d** Mutation of the E-box-like motif (CAACTTG) in the truncated *CrMYBR2* promoter (pro*CrMYBR2s*) affects its transactivation by HIVE. Schematic diagram showing the E-box and E-box-like motif in the full-length and two truncated *CrMYBR2* promoters, pro*CrMYBR2s* (-813 bp to -1 bp) and pro*CrMYBR2ss* (-347 bp to -1 bp). The E-box-like motif was mutated to CCCCCCG (m). **e** Vindoline content and relative expression of *CrMYBR2*, *T16H2*, *T3O*, *D4H,* and *DAT* in EV and *CrMYBR2*-overexpressing (*CrMYBR2*-OX) *C. roseus* seedlings. **f** Vindoline content and relative expression of *CrMYBR2*, *T16H2*, *T3O*, *D4H,* and *DAT* in EV and *CrIMYBR2*-VIGS leaves. **g** Mutation of the CR-box motif (m3, as indicated in Figure a) affects the activation of pro*D4H* by CrMYBR2. The effector and reporter (promoter-*GUS*) constructs were co-infiltrated into *C. roseus* seedlings. A plasmid containing the firefly *luciferase* (*LUC*) reporter was used as an internal control. GUS activity was normalized against luciferase activity. The control is the reporter with EV. Gene expression was measured by RT-qPCR. Alkaloids were extracted and analyzed by LC-MS/MS, and the concentrations of the alkaloids were estimated based on peak areas compared with standards. DW, dry weight. The data represent means ± SD from three biological replicates. Statistical significance was calculated using Student’s *t* test (* *p* <0.05 and ** *p* <0.01). **h** Yeast one-hybrid assay showing activation of the *D4H* promoter (pro*D4Hs*) by CrMYBR2. The plasmid harboring a GAL4 activation domain/CrMYBR2 fusion (pAD-CrMYBR2) was co-transformed with the pHIS2-pro*D4Hs* reporter plasmid. The yeast transformants were grown in either double selection medium (SD-Leu-Trp) or triple selection medium (SD-His-Leu-Trp) with 50 or 100 mM 3-AT.

A total of 10 CCA1-like and 8 R-R-type MYB-related TFs were identified in the *C. roseus* genome. Among them, two R-R-type factor genes (*CRO_T102972* and *CRO_T127742*) were co-expressed with *HIVE* and the genes associated with the vindoline pathway (Figure S2c). We designated *CRO_T102972* and *CRO_T127742* as *CrMYBR1* and *CrMYBR2*, respectively (Figure S2d). In *HIVE*-OX seedlings and *HIVE*-VIGS leaves, the expression of *CrMYBR2*, not *CrMYBR1*, was altered (Figure 4b and 4c). HIVE activated the full-length *CrMYBR2* promoter (pro*CrMYBR2,* 1,113 bp), increasing relative expression by 9-fold (Figure 4d). Eight E-box motifs were found in pro*CrMYBR2* (Figure 4d). Two truncated *CrMYBR2* promoters were generated, pro*CrMYBR2s* (excluding six E-boxes) and pro*CrMYBR2ss* (a further deletion to exclude an E-box-like element) (Figure 4d; Table S3). Transient promoter assays showed that HIVE had similar activity on pro*CrMYBR2s* and pro*CrMYBR2* (Figure 4d). However, the activation of pro*CrMYBR2ss* by HIVE decreased significantly compared to that of pro*CrMYBR2s* (Figure 4d). When the E-box-like motif (CAACTTG) was mutated to CCCCCCG (m) to generate the mutant promoter pro*CrMYBR2sm,* significant reduction of HIVE activation was resulted, similar to that of pro*CrMYBR2ss* (Figure 4d). The promoter regions of *T16H2*, *T3O*, *D4H,* and *DAT* do not contain E-box-like motifs.

The four genes associated with the vindoline pathway were upregulated by 3- to 5-fold when *CrMYBR2* was overexpressed in *C. roseus* seedlings (Figure 4e). The vindoline content increased significantly in *CrMYBR2*-OX seedlings relative to the EV control. When expression of *CrMYBR2* was repressed by VIGS in leaves (Figure 4f), transcript levels of *T16H2*, *D4H*, and *DAT* were reduced by 35%-50% compared to EV. Vindoline content was reduced by 55% in leaves of CrMYBR2-VIGS plants compared to EV. CrMYBR2 activated pro*D4H*-*GUS* by 3.5-fold. Mutation of the CR-box in the promoter (proD4Hm3) abolished the CrMYBR2 activation (Figure 4g). The interaction between CrMYBR2 and pro*D4Hs* was also confirmed in a yeast one-hybrid (Y1H) assay (Figure 4h).

#### bHLH repressor of vindoline-gene expression (BREVE), a repressor of vindoline biosynthesis

In Arabidopsis, bHLH proteins from subgroup Ia and IIIb are functionally associated (Kanaoka et al., 2008). A subgroup Ia bHLH gene (*CRO_T124980*; Figure S3) is co-expressed with *HIVE* and the genes associated with vindoline pathway (Table S1). Transient overexpression of *CRO_T124980* in *C. roseus* seedlings resulted in a 60-80% reduction in the expression of *T16H2*, *T3O*, *D4H*, and *DAT* (Figure 5a). Vindoline contents were also reduced significantly in seedlings overexpressing *CRO_T124980* relative to the EV control (Figure 5a). Knocking down *CRO_T124980* expression using VIGS in *C. roseus* leaves upregulated the expression of *T16H2*, *T3O*, *D4H*, and *DAT* by 2.5- to 3.5-fold (Figure 5b), and the vindoline content increased in leaves of *CRO_T124980*-VIGS plants compared to the EV plants (Figure 5b). Therefore, we termed CRO_T124980 “<u>b</u>HLH <u>re</u>pressor of <u>v</u>indoline-gene <u>e</u>xpression” (BREVE). BREVE repressed the promoter activities of all four genes associated with the vindoline pathway (Figure 5c). Repression of pro*D4H* activity by BREVE in the *Nicotiana benthamiana* leaf was abolished when the first E-box (m1) was mutated; however, mutation of the second E-box (m2) did not affect the repression (Figure 5d).

**Figure 5:**
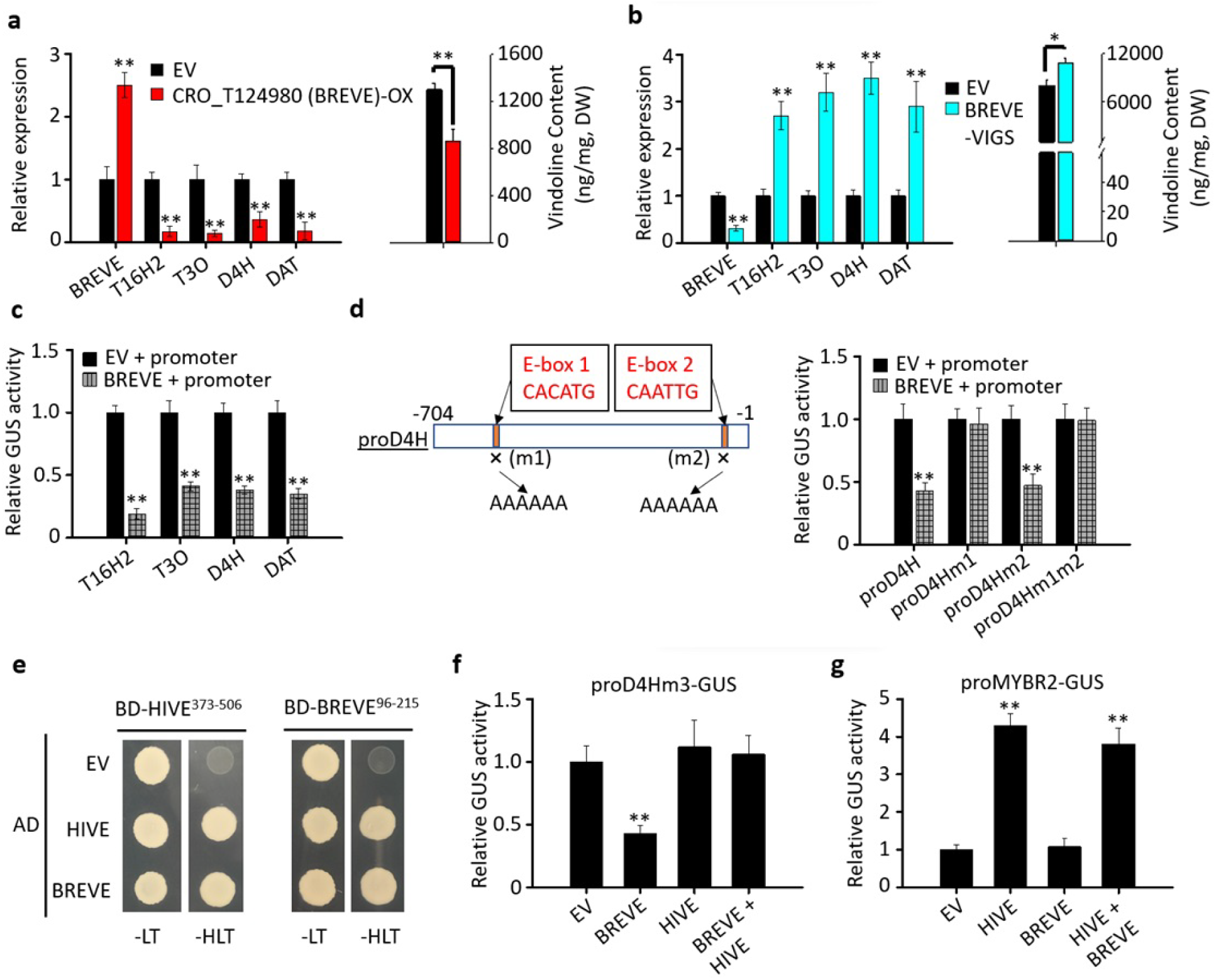
The repressive effect of BREVE on vindoline pathway genes is antagonized by HIVE. **a** Vindoline content and relative expression of *BREVE*, *T16H2*, *T3O*, *D4H,* and *DAT* in EV and *BREVE*-OX seedlings. **b** Vindoline content and relative expression of *BREVE*, *T16H2*, *T3O*, *D4H,* and *DAT* in EV and *BREVE*-VIGS leaves. Gene expression was measured by RT-qPCR. Alkaloids were extracted and analyzed by LC-MS/MS, and the concentrations of the alkaloids were estimated based on peak areas compared with standards. DW, dry weight. **c** Transactivation activities of the *T16H2*, *T3O*, *D4H*, and *DAT* promoters are repressed by BREVE. **d** Mutation of one E-box motif (E-box 2) affects the repression of the *D4H* promoter by BREVE. The E-box motifs in the *D4H* promoter were mutated to AAAAAA. **e** HIVE and BREVE form homo- and hetero-dimers in yeast cells. The bHLH domains of HIVE^373-506^ and BREVE^96-215^ were used as baits in the assay due to self-activation of the full-length proteins in yeast cells. Full-length HIVE and BREVE proteins were expressed as preys. Protein-protein interactions were detected by yeast growth on triple (-His-Leu-Trp) selection medium. Empty pAD-Gal4-2.1 vector (EV) was used as the negative control. **f** The repressive effect of BREVE on the *D4H* promoter is antagonized by HIVE. The CR-box mutant of the *D4H* promoter (pro*D4Hm3*, as described in Figure 3C) fused to the *GUS* gene was used as the reporter. BREVE and HIVE were used as effectors. **g** Transactivation of the *CrMYBR2* promoter by HIVE is not affected by BREVE. The *CrMYBR2* promoter fused to *GUS* gene was the reporter. HIVE and BREVE were used as effectors. Effector and reporter constructs were infiltrated in different combinations into *N. benthamiana* leaves. A plasmid containing the *LUC* reporter was used as the normalization control. LUC and GUS activities were measured 2 days after infiltration. GUS activity was normalized against LUC activity. The control was the reporter with EV. Values represent means ± SD from three biological replicates. Statistical significance was calculated using Student’s *t* test (* *p* <0.05 and ** *p* <0.01).

#### The repressive effect of BREVE is antagonized by HIVE

Yeast two-hybrid (Y2H) assays showed that HIVE interacts with BREVE, and that both HIVE and BREVE form homodimers through their bHLH domains (Figure 5e). BREVE, not HIVE, repressed the activity of pro*D4H* carrying the CR-box mutation (pro*D4Hm3*) (Figure 5f). However, pro*D4Hm3* activity was not affected when *HIVE* and *BREVE* were co-expressed (Figure 5f). In addition, while HIVE activated pro*CrMYBR2,* BREVE had no effect on pro*CrMYBR2,* and co-expression of *HIVE* with BREVE did not show reduction in pro*CrMYBR2* activation (Figure 5g). These results suggest that the HIVE-BREVE heterodimer antagonizes the repressive effects of BREVE on vindoline pathway genes but has little effect on activity of the *CrMYBR2* promoter.

#### HIVE/CrMYBR2/BREVE circuit regulates vindoline biosynthesis in response to herbivory and cold stress

Like *HIVE* and the vindoline pathway genes, the expression of *CrMYBR2* was not affected in DL but was induced in UPL after herbivory by *M. sexta,* while *BREVE* expression was not affected in DL or UPL (Figure 6a). *CrMYBR2* expression was downregulated after exposure to LT for 2 h, while the expression of *BREVE* remained unaffected by LT (Figure 6b). Expression of *CrMYBR2* and vindoline pathway genes was downregulated when seedlings overexpressing *HIVE* were exposed to LT (Figure 6c). At NT, however, overexpression of HIVE upregulates *CrMYBR2* and vindoline pathway genes (Figure 3a). This differential effects of HIVE on CrMYBR2 and vindoline pathway genes between NT and LT suggest that the HIVE specificity is affected by LT. When *C. roseus* seedlings overexpressing *CrMYBR2* or *BREVE* were exposed to LT, CrMYBR2 activated, but BREVE repressed the expression of the vindoline pathway genes (Figure 6d and 6e). This was similar to what was observed at NT (Figure 4e and Figure 5a). The results suggest that HIVE differentially regulates *CrMYBR2* and *BREVE* in response to temperatures and, together, the HIVE/CrMYBR2/BREVE regulatory circuit controls the differential accumulation of vindoline in response to herbivory and LT.

**Figure 6:**
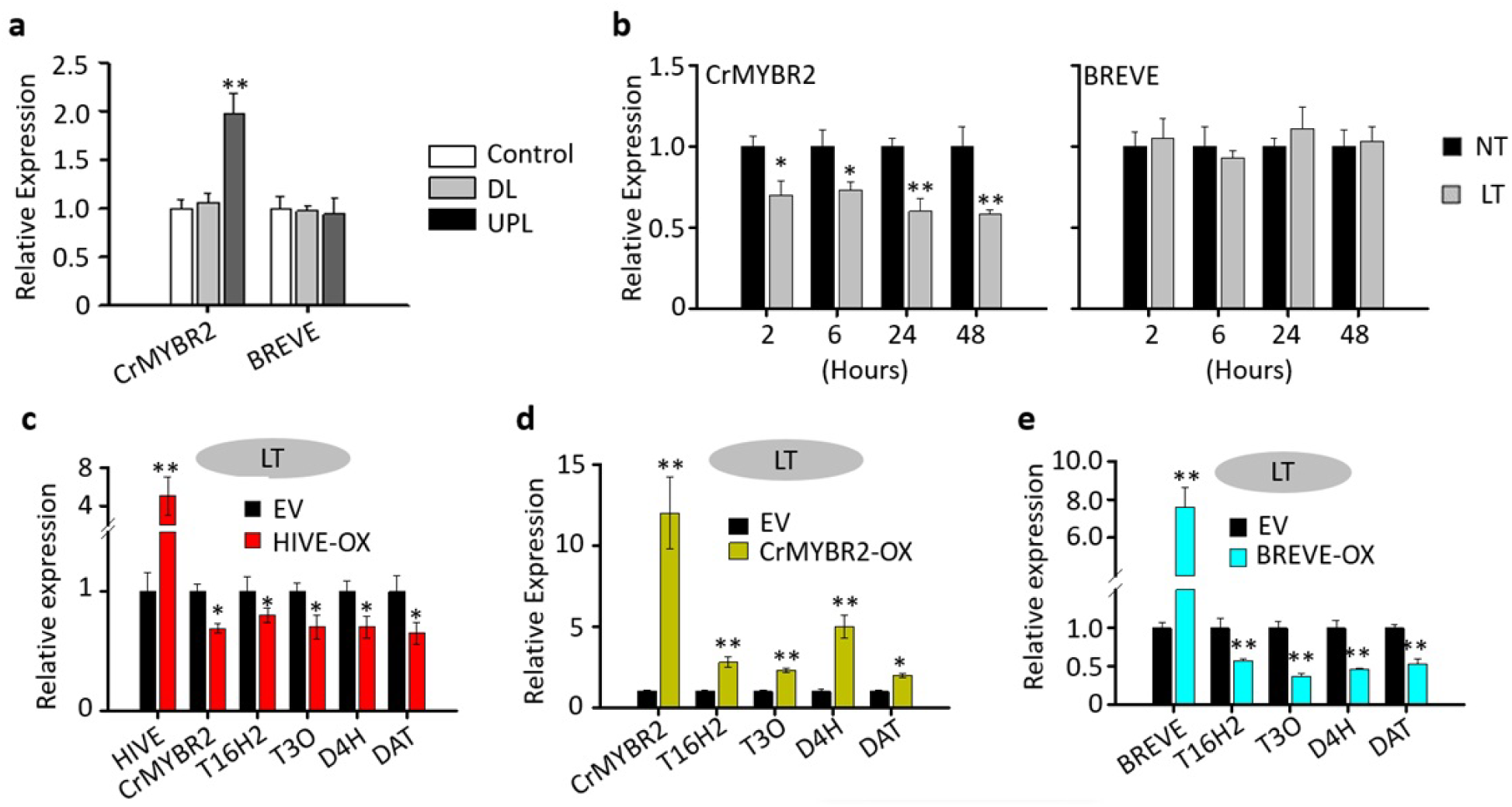
The HIVE-CrMYBR2 module responds to herbivory and cold stress to regulate vindoline biosynthesis. **a** Relative expression of *CrMYBR2* and *BREVE* in *C. roseus* leaves (control, DL, and UPL, as described in Figure 2d) after *M. sexta* herbivory. **b** Relative expression of *CrMYBR2* and *BREVE* in *C. roseus* seedlings exposed to LT (4℃) for 2, 6, 24, and 48 hours. **c** Relative expression of *HIVE*, *CrMYBR2,* and four vindoline pathway genes in EV and HIVE-overexpressing (*HIVE*-OX) *C. roseus* seedlings. **d** Relative expression of *CrMYBR2* and the vindoline pathway genes in EV and *CrMYBR2* overexpressing (*CrMYBR2*-OX) *C. roseus* seedlings. **e** Relative expression of *BREVE* and the vindoline pathway genes in EV and *BREVE* overexpressing (*BREVE*-OX) *C. roseus* seedlings. In **c**, **d**, and **e**, seedlings were exposed to LT for 2 hours before collection for RNA isolation. Gene transcript levels were measured by RT-qPCR. Data represent means ± SD of three biological replicates. Statistical significance was calculated using Student’s *t* test (* *p* <0.05 and ** *p* <0.01).

#### Phosphorylation alters HIVE specificity and suspends TIA biosynthesis under cold stress

Phosphorylation can change protein activities, and cell signaling in response to abiotic stresses in plants largely depends on the SnRK family protein kinases (Kulik, Wawer, Krzywińska, Bucholc, & Dobrowolska, 2011; Zhu, 2016). The SnRK2 subfamily kinases are involved in both specialized metabolism (Zhang et al., 2018) or cold tolerance (Ding et al., 2015). Seven SnRK2 family members (CrSnRK2.1 through CrSnRK2.7) were identified in the *C. roseus* genome, and CrSnRK2.7 is closely related to the Artemisia AaAPK1 (Figure S4a). Y2H assay show that HIVE interacts with CrSnRK2.7, but not CrSnRK2.6, another group III CrSnRK2 member (Figure 7a), suggesting that HIVE might be a substrate of CrSnRK2.7.

**Figure 7:**
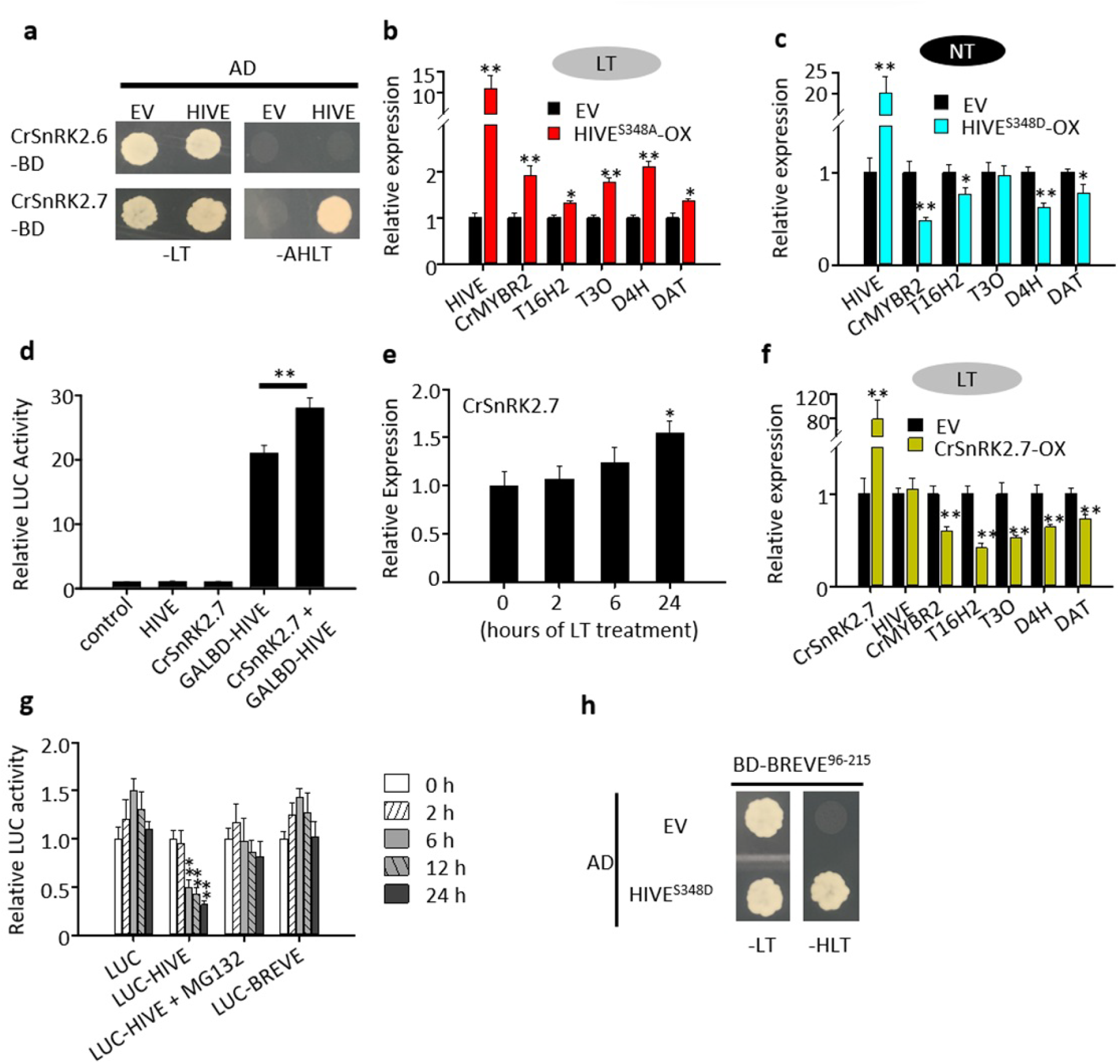
Phosphorylation alters HIVE transcriptional specificity in cold. **a** HIVE interacts with CrSnRK2.7 in yeast cells. *HIVE* was expressed as prey and *CrSnRK2.6* or *CrSnRK2.7* were expressed as baits in yeast cells. Protein-protein interactions were detected by yeast growth on quadruple (-His-Leu-Trp-Ade) selection medium. **b** Relative expression of *HIVE*, *CrMYBR2,* and the vindoline pathway genes in EV and *HIVE^S348A^*-overexpressing (*HIVE^S348A^*-OX) *C. roseus* seedlings exposed to LT (4℃) for 2 hours before sampling. HIVE^S348A^ is a non-phosphorylation mutant of HIVE. Gene expression was measured by RT-qPCR. **c** Relative expression of *HIVE*, *CrMYBR2,* and the vindoline pathway genes in EV and *HIVE*^S348D^-OX *C. roseus* seedlings at normal temperature (NT). HIVE^S348D^ is a phospho-mimic mutant of HIVE. **d** A protoplast-based assay demonstrates that CrSnRK2.7 enhances HIVE activity in plant cells. The GAL4 DNA binding domain (GALBD)-HIVE fusion protein activates the *GAL4*-promoter-driven *LUC* reporter in tobacco cells, and the activation was significantly increased when *GALBD-HIVE* was co-expressed with *CrSnRK2.7*. **e** Relative expression of *CrSnRK2.7* in leaves of two-month-old *C. roseus* plants exposed to LT for 2, 6, and 24 hours. **f** Relative expression of *CrSnRK2.7*, *HIVE*, *CrMYBR2,* and the vindoline pathway genes in EV and *CrSnRK2.7*-overexpressing (*CrSnRK2.7*-OX) *C. roseus* seedlings exposed to LT for 2 hours. **g** Degradation of HIVE in cold. HIVE or BREVE were translationally fused with the *luciferase* (*LUC*) reporter. The *LUC* control, *HIVE-LUC*, and *BREVE-LUC* were transiently overexpressed in *C. roseus* seedlings, and the seedlings were kept at LT for 0, 2, 6, 12, and 24 hours before sample collection. *C. roseus* seedlings were also treated with the proteasome inhibitor MG132 for 24 hours before exposure to LT. A plasmid containing the *GUS* reporter, driven by the *CaMV*35S promoter and *rbcS* terminator, was used as a normalization control. LUC activity was normalized against GUS activity. **h** HIVE^S348D^ interacts with BREVE in yeast cells. HIVE^S348D^ was expressed as prey in AH109 yeast cells. The bHLH domain of BREVE was used as bait. Protein-protein interactions were detected by yeast growth on triple (-His-Leu-Trp) selection medium. In **c** and **h**, the empty pAD-Gal4-2.1 vector (EV) was the negative control. Data represent means ± SD of three biological replicates. Statistical significance was calculated using Student’s *t* test (* *p* <0.05 and ** *p* <0.01).

SnRK2 family kinases phosphorylate serine (S) in two common motifs: SD and RxxS (x represents any amino acid) (Wang et al., 2013). Four putative SnRK2 targeted phosphor motifs were found in HIVE, among which S348 was investigated due to its conservation to a known phosphorylation site in the subgroup IIIb bHLH (Ding et al., 2015). Two HIVE mutants, HIVE^S348A^ and HIVE^S348D^, were generated, in which S348 was substituted with either alanine (A) to generate phosphor-null or an aspartate (D) to generate a phospho-mimic form of HIVE. When *HIVE^S348A^* was transiently overexpressed in *C. roseus* seedlings and then exposed to LT, the expression of *CrMYBR2* and genes associated with the vindoline pathway was upregulated (Figure 7b). Overexpression of *HIVE^S348D^* resulted in a decrease in the relative expression of *CrMYBR2* and the genes associated with the vindoline pathway at NT (Figure 7c).

A fusion protein consisting of the GAL4 DNA binding domain fused in frame to HIVE (GALBD-HIVE) activated the *GAL4*-promoter-driven *LUC* reporter gene in tobacco cells, and the activation was significantly increased when *GALBD-HIVE* was co-expressed with *CrSnRK2.7* (Figure 7d). *CrSnRK2.7* transcription increased by ∼50% after 24 h of LT treatment (Figure 7e). At LT, transient overexpression of *CrSnRK2.7* in *C. roseus* seedlings had no effect on the expression of *HIVE* but repressed the expression of *CrMYBR2* and the vindoline pathway genes (Figure 7f).

#### HIVE, not BREVE, is degraded by the 26S/UPS proteasome

To determine whether the stability of HIVE is affected by cold, we fused the firefly *LUC* reporter with *HIVE* (*LUC-HIVE*) and transiently overexpressed the fusion gene in *C. roseus* seedlings. LUC activities in seedlings infiltrated with the *LUC-HIVE* construct were significantly reduced after 6 h of LT treatment compared to the control (Figure 7g). LUC activities in seedlings treated with the proteosome inhibitor MG132 were similar to those of the control, suggesting that HIVE is degraded at LT through 26S/UPS (Figure 7g). Unlike HIVE, BREVE appeared to be not degraded at LT as the LUC-BREVE activities remained unchanged (Figure 7g). We have revealed that at NT, HIVE dimerizes with BREVE, thus preventing BREVE from repressing vindoline biosynthesis (Figure 5f). A Y2H assay showed that the phospho-mimic HIVE^S348D^ retained its interaction with BREVE (Figure 7h), indicating that HIVE-BREVE dimerization is not affected by phosphorylation. The degradation of HIVE likely results in breakdown of the HIVE/BREVE heterodimer.

#### HIVE signals cold tolerance at a low temperature

The C-repeat binding factors (CBFs) are key components of the cold signaling pathway in plants. Genes encoding three CBF-like proteins, CrCBF1, CrCBF2, and CrCBF3, were identified in the *C. roseus* genome. Only *CrCBF1* expression was induced by LT (Figure 8a). While HIVE induced CrCBF1 transcription at both NT and LT, the induction level was significantly higher at LT than NT (Figure 8b). Compared to EV, overexpression of *HIVE* and CrCBF1 led to significant decreases in relative electrolyte leakage (REL) and the malondialdehyde (MDA) contents (Figure 8c and 8d), confirming the positive roles of HIVE and CrCBF1 in cold tolerance. Overexpression of *HIVE^S348A^* in LT-treated *C. roseus* seedlings caused a significant reduction in *CrCBF1* expression (Figure 8e). In contrast, overexpression of *HIVE^S348D^* increased the expression of *CrCBF*1 at NT by ∼3.5-fold (Figure 8f). Additionally, overexpression of *CrSnRK2.7* in LT-treated *C. roseus* seedlings significantly induced *CrCBF1* expression compared to EV (Figure 8g). In tobacco cells, HIVE activated the *CrCBF1* promoter at LT, but not at NT (Figure 8h, right panel). *Cis*-element analysis of the *CrCBF1* promoter revealed two adjacent bHLH TF binding motifs, one E-box (CAACTG) and one E-box-like (CAACTAG), in the CrCBF1 promoter (Figure 8h, left panel). Mutation of the E-box-like motif eliminated the CrCBF1 promoter activation by HIVE (Figure 8h, right panel).

**Figure 8:**
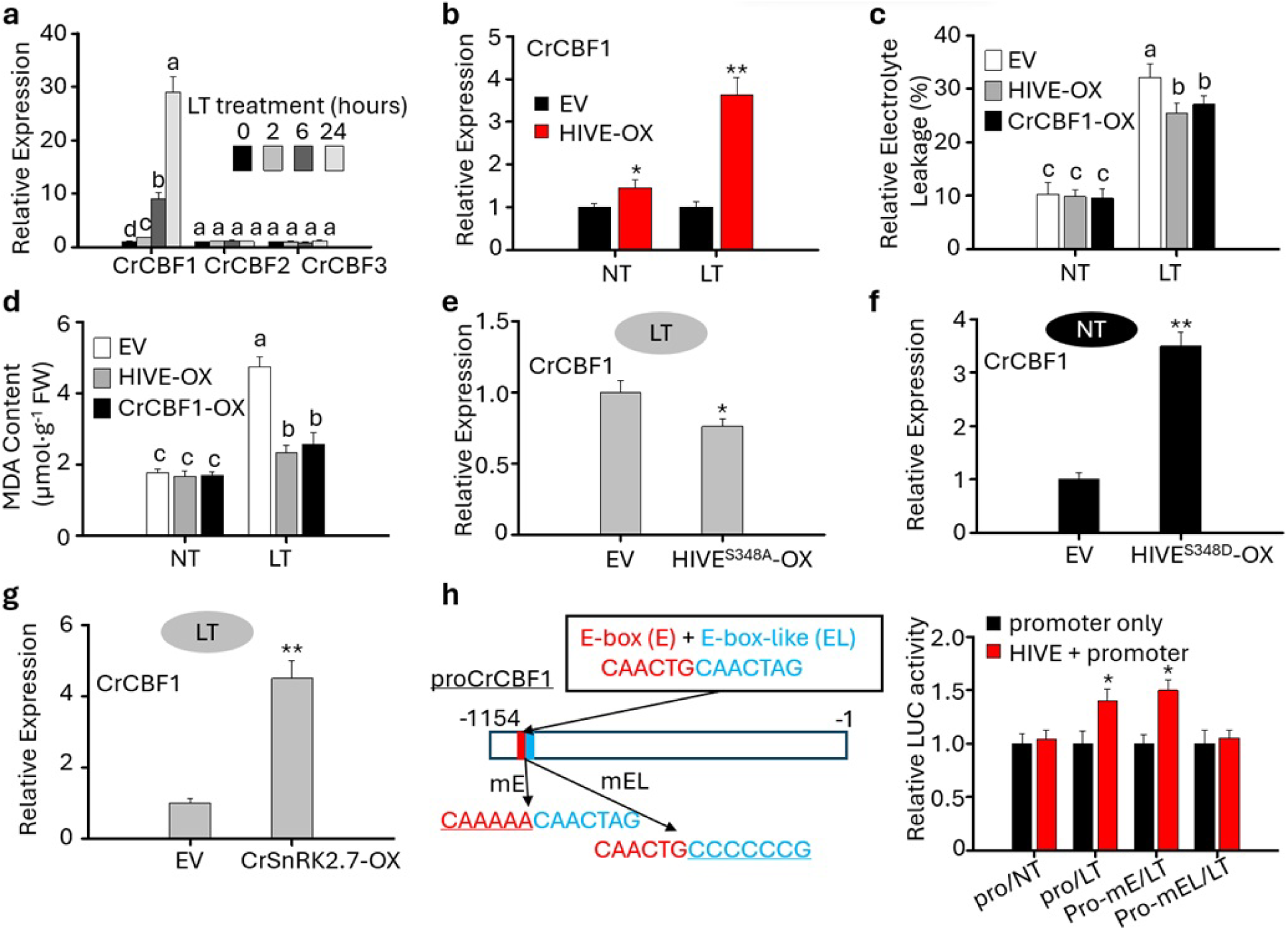
HIVE signals cold tolerance at a low temperature. **a** Relative expression of three *CBF*-like genes (*CrCBF1*, *CrCBF2,* and *CrCBF3*) in leaves of two-month-old *C. roseus* plants exposed to LT (4℃) for 2, 6, and 24 hours. Gene expression was measured by RT-qPCR, and the *C. roseus RPS9* gene was used as an internal reference gene. **b** Relative expression of *CrCBF1* in EV and *HIVE*-OX *C. roseus* seedlings at NT or exposed to LT for 2 hours. Relative electrolyte leakage **(c)** and MDA contents **(d)** in EV control, HIVE-OX and CrCBF1-OX *C. roseus* seedlings after kept in NT or exposed to LT (4℃) for 2 days. **e** Relative expression of *CrCBF1* in EV and *HIVE^S348A^*-OX *C. roseus* seedlings exposed to LT for 2 hours. **f** Relative expression of *CrCBF1* in EV and *HIVE^S348D^*-OX *C. roseus* seedlings. **g** Relative expression of *CrCBF1* in EV and *CrSnRK2.7*-OX *C. roseus* seedlings exposed to LT for 2 hours. **h** HIVE transactivates the *CrCBF1* promoter (pro*CrCBF1*) through an E-box-like (CAACTAG) motif at LT in tobacco cells. Left panel, a schematic diagram showing the overlapping E-box and E-box-like motifs in the *CrCBF1* promoter. E-box motif was mutated to CAAAAA (mE). E-box-like motif was mutated to CCCCCCG (mEL). A plasmid containing the GUS reporter was used as a normalization control. LUC activity was normalized against GUS activity. Data represent means ± SD of three biological samples. In **a**, **c** and **d**, different letters denote statistical differences as assessed by one-way ANOVA and Tukey’s honestly significant difference test (P < 0.05). In **b**, **e**, **f**, **g** and **h**, statistical significance was calculated using the Student’s t test (* P < 0.05; ** P < 0.01).

## Discussion

### “Timebomb” - a compartmentalized plant defense against herbivory with minimal fitness costs

Both plants and their herbivore pests have evolved various strategies to avoid each other’s defense systems during the coevolutionary arms race. Plants employ efficient biochemical defensive systems, such as specialized metabolites, that are critical to overcoming intense herbivore attacks (Ehrlich & Raven, 1964; Mithöfer & Boland, 2012; Negin & Jander, 2023; Speed, Fenton, Jones, Ruxton, & Brockhurst, 2015; War et al., 2012). Our results revealed a TIA “timebomb” in *C. roseus* leaves that effectively influences the survival of a chewing insect, *M. sexta,* as a resistance-proof and sustainable chemical defensive strategy against herbivory (Figure 1). Vindoline and catharanthine are either stored in separate compartments or accumulated in specific cells without sufficient dimerization. Tests of these two TIAs individually showed mild/no toxicity against *M. sexta,* whereas the mixture of the two was highly toxic, showing a clear synergistic effect (Figure 1a, Table 1). The selection pressure for resistance correlates with toxicity: usually the more toxic the defensive chemicals, the stronger the selection pressure (Hawkins, Bass, Dixon, & Neve, 2019; South & Hastings, 2018). The negligible toxicity of the individual TIAs suggests that there is no selection pressure for herbivores feeding on *C. roseus* to develop resistance against each TIA. When vindoline and catharanthine are consumed together, the high toxicity would likely annihilate the herbivores, providing no chance to establish a viable population or resistance. This two-component “timebomb” would be more effective against chewing insects compared to piercing insects, as the latter is less likely to be exposed to both precursors (or enzymes and substrates) simultaneously. Documented natural pests of *C. roseus* are, indeed, limited to arthropods with piercing-sucking mouthparts, including aphids (Maciel et al., 2011; Samad et al., 2008), whiteflies (Francis et al., 2016), scale insects (Kondo, Amalia Ramos-Portilla, Peronti, & Gullan, 2016), the pink hibiscus mealybug (Chong, Aristizábal, & Arthurs, 2015), and two-spotted spider mite (Haque, Islam, Naher, & Haque, 2011).

The mode of action (MOA) of the TIA “timebomb” also suggests an induced qualitative defense mechanism. This mechanism may be favored by natural selection as a sustainable strategy due to its minimal fitness cost. The high toxicity can only be achieved by chewing the leaves, thus accessing the toxic mixture of vindoline and catharanthine. After consumption of *C. roseus* leaf tissue, the ratio of (anhydro)vinblastine to catharanthine and vindoline was dramatically increased in the larval gut of *M. sexta* compared to the amounts measured in the whole leaf (Dugé de Bernonville et al., 2017). Biosynthesis of anhydrovinblastine is costly for the plant, requiring energy and resources (Cipollini, Walters, & Voelckel, 2018; Sottomayor & Ros Barceló, 2003). The “timebomb”, however, is not active until the plant is attacked by chewing herbivores. In this case, carrying non-toxic (inactive) precursors other than constitutively accumulating the highly toxic (active) dimers would benefit *C. roseus* by minimizing the fitness costs associated with host-derived chemical defenses. In addition, toxicity of the dimeric mixture is evident even at a lower dose, suggesting a qualitative mechanism in which chemical defense can be achieved at lower concentrations and requires a minimal tradeoff with fitness costs (Coley, Bryant, & Chapin, 1985).

### Crosstalk between herbivory resistance and cold tolerance in *C. roseus*

The influences of abiotic and biotic stresses on various developmental and metabolic processes in plants have been studied for many years; however, crosstalk between abiotic and biotic stresses that alters plant responses in ever-changing environments are not well understood (Bostock, Pye, & Roubtsova, 2014; Fujita et al., 2006). In recent years, increasing attention has been paid to the interplay and the impacts of different signals that generate defense responses against a plethora of environmental cues (Desaint et al., 2021; Gu et al., 2023; Suzuki, Rivero, Shulaev, Blumwald, & Mittler, 2014). Our finding provides a prime example of how plants prioritize their efforts to manage multiple stresses concurrently. Haller et al. (Haller et al., 2020) showed that salt stress in Arabidopsis increases susceptibility to *Botrytis cinerea,* a necrotrophic fungus. We showed that vindoline biosynthesis is induced by herbivory but repressed by cold stress (Figure 2). *C. roseus* produces vindoline and catharanthine, the two constituents of the “timebomb” at normal temperature. Insect activities are reduced when temperature decreases. *C. roseus* switches metabolic efforts to tolerate cold stress, indicating potential crosstalk between cold stress and herbivory in the regulation of vindoline biosynthesis. The differential responses to herbivore defense and cold tolerance in *C. roseus* represent a balancing strategy to minimize the energy cost when responding to environmental stresses.

### Transcriptional reprogramming of specialized metabolism allows plants to fine-tune their stress responses to accommodate multiple stressors

Our combined work demonstrated that HIVE is a dual-function regulator fine-tuning the needs for both chemical defense and cold tolerance in *C. roseus*. At a normal temperature, HIVE triggers TIA biosynthesis through activation of CrMYBR2 in response to herbivory. At a low temperature, however, the phosphorylated HIVE represses *CrMYBR2* while activating the CBF-mediated cold tolerance pathway, which is conserved in non-cold-acclimated *C. roseus*. In cold-acclimated *Arabidopsis*, the CBF-mediated cold tolerance pathway is, in part, regulated by the bHLH TF ICE1 (Chinnusamy et al., 2003). However, ICE1 (a significantly smaller protein with low sequence identity outside of the bHLH domain compared to HIVE) is not involved in the biosynthesis of specialized metabolites or chemical defense.

The disruption of plant metabolism by stresses is often mediated through crosstalk that involves phytohormones, TFs, and signaling cascades. It is thus reasonable to envision that a transcriptional hub coordinates the balance between two diverse types of stress. Here, the crosstalk between chemical defense and cold tolerance in *C. roseus* was shown to be affected by several TFs, including HIVE, CrMYBR2, and BREVE (Figure 9). At normal temperatures and in response to biotic stress, HIVE activates *CrMYBR2,* which in turn induces the expression of the vindoline biosynthetic pathway genes, resulting in increased leaf accumulation of vindoline (Figure 3 and 4), a key component of the “timebomb”. In addition to CrMYBR2 activation, HIVE also forms heterodiminer to sequester the repressor BREVE (Figure 5e and 5f), which is not induced by herbivory (Figure 6a). As such, the net outcomes of herbivore-induced *HIVE* expression result in increased vindoline and TIA production. Exposure to low temperature induces *CrSnRK2.7*, hence increasing the level of HIVE phosphorylation. Phosphorylated HIVE represses *CrMYBR2* (Figure 7b and 7f). Upon prolonged exposure to cold, HIVE is degraded by the 26S/UPS proteasome (Figure 7g), leading to the release of BREVE which strongly represses expression of the vindoline pathway genes. Under cold conditions, therefore, HIVE rapidly switches its specificity through phosphorylation, resulting in a decrease in vindoline production. The HIVE-centric TF circuit coordinates the production of toxic specialized metabolites in response to herbivory and activation of the cold-signaling pathway at low temperatures. Our findings are in accord with a recent study showing how the brassinosteroid (BR)-induced TF BES1 coordinates growth and Ultraviolet-B (UV-B) stress response in Arabidopsis. BES1 promotes growth and inhibits flavonoid biosynthesis by directly repressing the transcription of key flavonoid regulator genes, such as *MYB11*, *MYB12*, and *MYB111*. In order to manage the detrimental effects of UV-B exposure, however, *BES1* expression is down-regulated to promote the biosynthesis of UV-absorbing flavonoid compounds (Liang et al., 2020).

**Figure 9:**
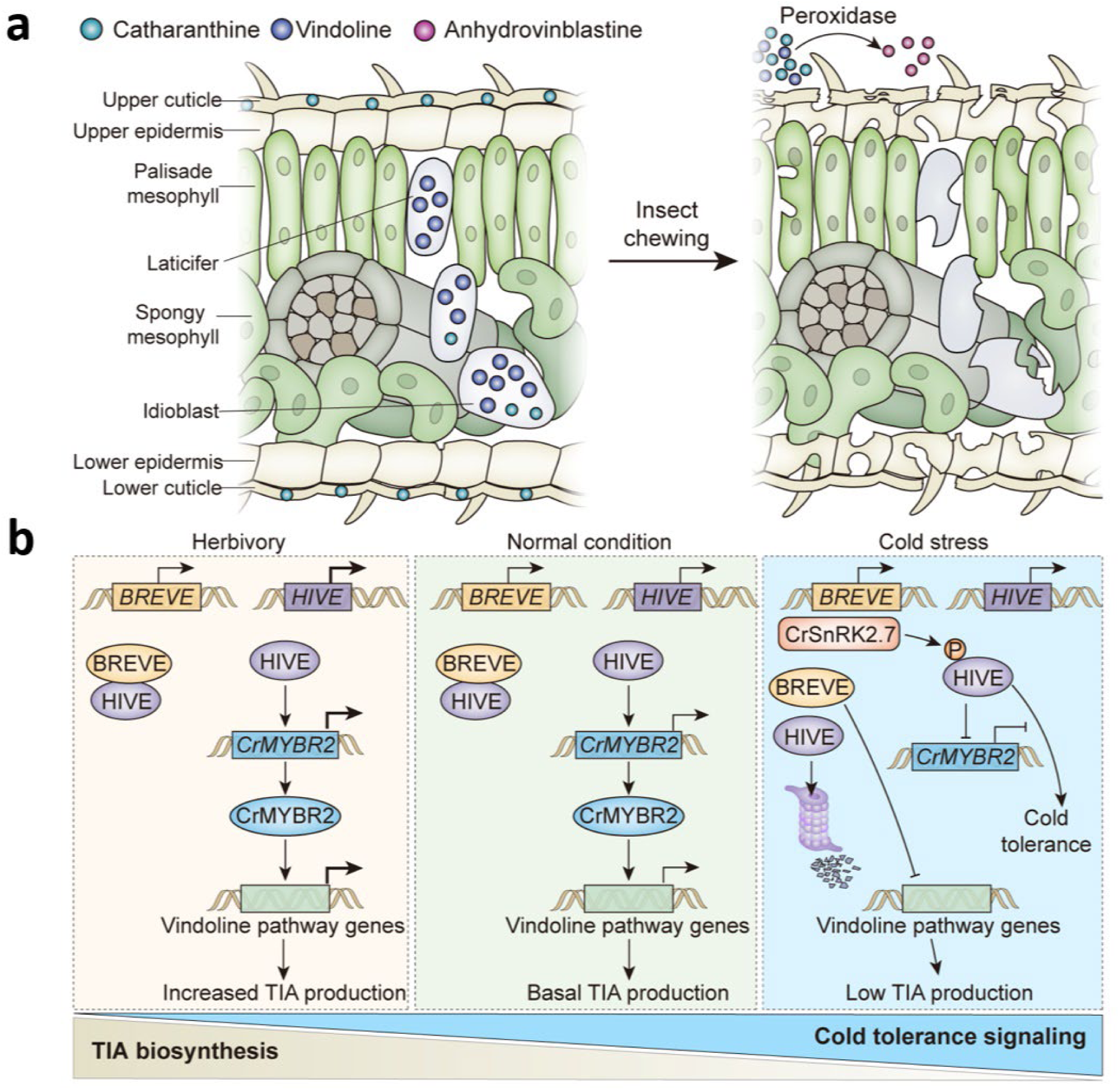
Transcriptional reprogramming of terpenoid indole alkaloid biosynthesis in response to herbivory- and cold-stresses in *C. roseus*. **a** “Timebomb” strategy involves vindoline and catharanthine condensation to defend against *Manduca sexta* feeding. Catharanthine is localized at both the leaf surface and the laticifer and idioblast cells, while vindoline accumulates exclusively in laticifer and idioblast cells. Leaf cells are disrupted by chewing herbivores leading to the dimerization of catharanthine and vindoline, possibly by peroxidase (PRX1). **b** Left panel, at normal temperatures when the herbivory pressure is high (compared to normal condition, middle panel), *HIVE* and *CrMYBR2* are induced and the HIVE-CrMYBR2 cascade positively regulates vindoline biosynthesis. HIVE interacts with and sequesters BREVE, thus antagonizing the repressive effect of BREVE on vindoline biosynthesis. The net transcriptional output favors TIA biosynthesis. Right panel, at low temperatures, TIA biosynthesis is suppressed while cold tolerance signaling is activated. CrSnRK2.7 is activated to phosphorylate HIVE. Phosphorylated HIVE is still capable of interacting with and sequestering BREVE. Phosphorylated HIVE represses *CrMYBR2* expression but preferentially activates cold tolerance pathway. With prolonged exposure to cold, HIVE is degraded by the 26S/ubiquitin proteasome, thus releasing BREVE which strongly represses the vindoline biosynthesis genes.

### Summary and perspectives

In this study, we investigated a mechanism by which plants prioritize their defensive strategies to manage the herbivory and cold stresses concurrently. Our combined results support the overarching hypothesis that HIVE-mediated transcriptional reprogramming of plant specialized metabolism can fine-tune stress management in *C. roseus.* Firstly, we documented “timebomb”, a unique insecticidal mode of action in *C. roseus*. Given that the TIA “timebomb” can only be activated when the two spatially isolated, non-toxic monomeric precursors are dimerized by herbivory, this qualitative chemical defense mechanism against herbivory is not only adaptive and innovative, but also sustainable. Without herbivory, the presence of insecticidal dimer is negligible, suggesting that no/minimal fitness cost is associated with the TIA “timebomb”.

Secondly, we identified a HIVE-centric, multifactorial regulatory circuit in *C. roseus* that strategically modulates the availability of TIAs in response to herbivory and cold stress. Specifically, at a normal temperature, HIVE initiates vindoline biosynthesis by activating a transcriptional activator (*CrMYBR2*) and sequestering a repressor (*BREVE*), leading to the basal accumulation of vindoline. Upon herbivory, *HIVE* expression is significantly induced, resulting in an upregulation of *CrMYBR2* and vindoline pathway genes. However. at a lower temperature, cold-triggered phosphorylation by CrSnRK2 switches HIVE to serve as a repressor of CrMYBR2 and an enhancer of CBF-mediated cold tolerance pathway. Furthermore, phosphorylation leads to the proteasomal degradation of HIVE, resulting in the release of BREVE that directly represses the vindoline pathway. In summary, HIVE-mediated transcriptional reprogramming allows this herbaceous perennial to allocate their metabolic resources for chemical defense at a normal temperature (25°C), when herbivory pressure is high, but switches to cold tolerance under a cooler temperature (4°C), when insect infestation is secondary. Finally, it is tempting to speculate that a transcriptional reprogramming by HIVE-like transcription factor(s) to strategically managing multiple stressors concomitantly might be a rule rather than an exception in plants.

## Materials and Methods

### Mode of action of the TIA “timebomb”, a qualitative chemical defense in *C. roseus*

#### Insect rearing

Young tobacco plants (*N. tabacum* cv. ‘Samsun NN’) were grown in individual pots at 27°C in a greenhouse and fresh leaves were used as hatching and feeding substrates for larvae of the tobacco hornworm, *M. sexta*, (eggs obtained from Carolina Biological Supply Company, NC, US). All newly hatched *M. sexta* larvae were collected and reared in Petri dishes (d = 3.5 cm) with tobacco leaves prior to the feeding experiment (60% RH, 27°C).

#### Feeding bioassay

The effect of catharanthine and vindoline on *M. sexta* larvae was examined by feeding the larvae on tobacco leaves supplied with the *Catharanthus* alkaloids. Agar plates with 2 ml of 1.5% (w/w) agar applied to the bottom of the Petri dishes (d = 3.5 cm) were used as the testing arenas (Figure 1c). Leaf disks (d = 15 mm) were punched out of tobacco leaves. Solutions of both alkaloids at a concentration of 2.19 μmol/ml were prepared in methanol, then serially diluted to yield solutions of 1-, 0.1-, and 0.01-fold concentrations (Table S4). Alkaloid solutions were carefully and evenly applied to the adaxial surfaces of the leaf disks. To test the dietary toxicity of the TIAs, three testing groups were conducted simultaneously: the two individual TIA compounds and the 1:1 ratio mixture at the same concentration. Alkaloid solution (40 μL) was applied to the leaf surfaces. Based on the parameters (including the treated number of alkaloids, the fresh weight of *C. roseus* and tobacco leaf disks, and the alkaloid contents in *C. roseus*), treatment with the 10-fold dilution (0.219 μmol/ml) is approximately equivalent to the actual alkaloid contents in fresh *C. roseus* leaves. The same volume of methanol was used as the control. All treated leaf disks were placed in a fume hood to dry for half an hour before being transferred onto the agar plate. A newly hatched *M. sexta* larva (within 24 h) was then introduced onto the leaf disk. Petri-dishes were kept in a greenhouse during the bioassay. The mortality and body weights of the larvae were documented at 24, 48, 72. and 96 h. Images of living larvae were taken daily to document the changes in the *M. sexta* larvae due to the influence of the TIAs. The area of each leaf disk was measured before and after the survey (either when the larvae were dead or when the assay ended at 96 h), then measured using Fiji software(Schindelin et al., 2012) to estimate the amount of leaf tissue consumed. *M. sexta* larvae were considered dead if no movement was observed when stimulated with a small brush. Three replicates were conducted for each treatment with ten individuals in each replicate. Five leaf disks were used in each replicate.

#### Synergy analysis

The effect of the alkaloid mixture on mortality was tested by comparing the observed to the expected mortalities following the methods in Hummelbrunner and Isman (Hummelbrunner & Isman, 2001) and Pavela (Pavela, 2015). Expected mortalities were calculated using the following formula: *E* = *O*_1_ + *O*_2_ (1 – *O*_1_), where *E* is the expected mortality and *O*_1_ and *O*_2_ are the observed mortalities for vindoline and catharanthine at each concentration. Chi-square values were used to compare expected with observed to designate the mixed effects as either antagonistic/synergistic or additive: *χ*^2^ = (*O*_m_ - *E*)^2^/*E*, where *O*_m_ is the observed mortality of the mixtures and *E* is the expected mortality. The mixed effect was considered to be synergistic when the observed mortality was higher than the expected and *χ*^2^ > 3.84, while it was considered to be additive when *χ*^2^ < 3.84 (*χ*^2^ = 3.84 with *df* = 1 and *p* = 0.05).

### Crosstalk between chemical defense and cold tolerance in *C. roseus*

#### Catharanthus roseus

Seeds of *C. roseus* (L.) G. Don cultivar ‘Little Bright Eye’(NESeed, USA) were surface sterilized and germinated on half-strength Murashige-Skoog (MS) salts medium(Patra, Pattanaik, Schluttenhofer, & Yuan, 2018). To examine the effect of low temperature on TIA biosynthesis, TIA contents and the expression of selected genes associated with the vindoline pathway were measured in 10-day-old seedlings or three-week-old plants with their first pair of true leaves.

#### The effects of low temperature and herbivory on TIA biosynthesis, individually

Plants were treated with low temperature (4℃) for 2, 6, 24, and 48 hours. Aerial parts of the seedlings or the first pair of true leaves of plants were collected for RNA isolation and alkaloid extraction. To determine the effects of herbivory-induced stress on vindoline biosynthesis, third instar *M. sexta* larvae were placed on leaves of eight-week-old *C. roseus* plants for two days. Vindoline content and the expression of genes associated with the vindoline pathway were measured in DL and UPL. *IPRT1*, a gene associated with biotic defense signaling, was used as a marker for the herbivory response (Dugé de Bernonville et al., 2017).

#### Crosstalk between low temperature- and herbivory-induced stress in the regulation of vindoline biosynthesis

To examine the crosstalk between LT- and herbivory-induced stress in the regulation of vindoline biosynthesis, the effect of LT on herbivory-induced vindoline biosynthesis was examined. After feeding by *M. sexta* for two days at NT, the treated *C. roseus* and control plants were immediately transferred to LT for two days. DL and UPL were collected for RNA isolation and alkaloid extraction. The sequences of primers used in this study are listed in Table S5. The concentrations of the alkaloids were calculated using standard curves.

### Transcriptional reprogramming of TIA biosynthesis in response to herbivory- and cold-stresses

#### Identification of novel regulators of vindoline biosynthesis

To identify novel regulators of the vindoline pathway, we focused on the bHLH TF family because; (1) E-box motifs, binding sites for bHLH TFs, are present in the promoters of four cold/herbivory-responsive vindoline pathway genes, and (2) bHLH TFs are known regulators of both plant specialized metabolism and cold tolerance (Chinnusamy et al., 2003; Goossens, Mertens, & Goossens, 2017; Patra et al., 2018; Xie et al., 2012). Transcriptomic resources of C. roseus (accession no. SRA030483) were used to perform co-expression analysis of bHLH TFs and the four genes associated with the vindoline pathway. Sequences of all genes were downloaded from the latest version of the C. roseus genome from Dryad Digital Repository. A BLAST search was performed to identify putative TF genes in the genome. To further validate the BLAST results, phylogenetic trees were constructed and visualized using the Neighbor-Joining (N-J) method with MEGA5.1 software. The statistical reliability of the individual nodes of the newly constructed trees was assessed by bootstrap analyses with 1,000 replications (Altschul et al., 1997). Amino acid alignments were performed using the MAFFT method (Katoh & Standley, 2013). To analyze the expression patterns of bHLH TF genes and selected genes in the vindoline pathway in C. roseus, transcriptomes from five different tissues (seedling, immature leaf, mature leaf, stem, and root; accession number: SRA030483) were used. Hierarchical clustering was performed (Paul et al., 2017).

#### Plasmid construction, transient overexpression in C. roseus seedlings

To elucidate the role of selected TFs in vindoline biosynthesis, the genes were transiently overexpressed in *C. roseus* seedlings and knocked-down in *C. roseus* leaves using virus-induced gene silencing (VIGS). For transient overexpression, TFs were cloned into a modified pCAMBIA1300 vector containing the *CaMV*35S promoter and the *rbcS* terminator. Mutants were generated by PCR-based site-directed mutagenesis (Pattanaik, Werkman, Kong, & Yuan, 2010). Seedlings of *C. roseus* were transiently transformed with the plasmids using the FAST method with some modifications (Weaver, Goklany, Rizvi, Cram, & Lee-Parsons, 2014). Briefly, *Agrobacterium tumefaciens* GV3101 harboring the plasmid was grown on Luria-Bertani (LB) plates containing 100 µg ml^−1^ kanamycin, 50 µg ml^−1^ gentamicin and 30 µg ml^−1^ rifampicin. A single colony was transferred to 2 ml liquid LB medium containing the same antibiotics and incubated at 250 rpm and 28°C overnight. The *Agrobacterium* cells were then sub-cultured in 20 ml liquid LB medium for 16 h at 250 rpm and 28°C. The *Agrobacterium* cultures were then centrifuged, and the pellet was resuspended in infiltration buffer (10 mM MgCl_2_, 10 mM MES, 100 µM acetosyringone) at an OD_600_ of 1.0, followed by incubation at 28°C for at least 3 h. Seven-day-old *C. roseus* seedlings were immersed in the *Agrobacterium* cultures for 1 h. Following vacuum infiltration, seedlings were washed five times with sterile distilled water and lain on sterile wet filter papers in Petri dishes. After three days of incubation at NT, seedlings were either directly collected or transferred to a cold room (4℃) for two hours. The aerial parts of the seedlings were collected for RNA extraction and alkaloid analyses.

#### Plasmid construction and promoter assays

To determine whether the selected TFs directly activate the genes associated with the vindoline pathway, the promoter regions of these genes were individually cloned upstream of the *GUS* reporter and were either transiently transformed into *N. benthamiana* leaves or *C. roseus* seedlings alone or with a TF. The reporter plasmids for promoter assays were generated by replacing the *CaMV*35S promoter in a modified pKYLX71 vector containing the *GUS* reporter and *rbcS* terminator with vindoline pathway gene (*T16H2*, *T3O*, *D4H* and *DAT*) promoters (Liu, Patra, Pattanaik, Wang, & Yuan, 2019). In addition, the promoter of *CrMYBR2* was cloned here. Mutants of the E-box or CR-box elements in the *D4H* promoter, and E-box-like in the *CrMYBR2* promoter, were generated by site-directed mutagenesis (Pattanaik et al., 2010). pCAMBIA1300 vectors containing *HIVE*, *BREVE* or *CrMYBR2* were used as the effector plasmids. The LUC reporter driven by *CaMV*35S promoter and *rbcS* terminator was used as an internal control in the leaf infiltration assays. Infiltration solutions were prepared as described in the FAST method. Before infiltration, effector, reporter and internal control solutions were combined at 1:1:1 ratios and mixed well. Infiltration of *N. benthamiana* leaves and transient transformation of *C. roseus* seedlings were performed as previously described (Kumar & Bhatia, 2016; Weaver et al., 2014). Two days after infiltration, LUC and GUS activities were measured in the leaf discs as previously described (Pattanaik et al., 2010).

#### Transient tobacco protoplast assays

For transient protoplast assays, the reporter plasmids were generated by cloning *CrCBF1* promoter in a modified pUC vector containing the LUC reporter and *rbcS* terminator. Mutant of the E-box, E-box-like in *CrCBF1* promoter was generated by site-directed mutagenesis. The effector plasmids were constructed by cloning *CrSnRK2.7* or *CrHIVE* into a modified pBS vector under the control of the *CaMV*35S promoter and *rbcS* terminator. For protoplast-based activity assay, CrHIVE were fused to the GAL4 DNA binding domain in a pBS plasmid containing *mirabilis mosaic virus* (*MMV*) promoter and *rbcS* terminator. The *GUS* reporter, driven by *CaMV*35S promoter and *rbcS* terminator, was used as an internal control in the protoplast assay. Protoplast isolation from *Nicotiana tabacum* cell suspension cultures and electroporation with supercoiled plasmid DNA were performed using previously described protocols (Pattanaik et al., 2010). The reporter, effector and internal control plasmids were electroporated into tobacco protoplasts in different combinations; luciferase and GUS activities in transfected protoplasts were measured as described previously (Pattanaik et al., 2010).

#### Protein degradation assay in C. roseus seedlings and Cold tolerance assay of C. roseus seedlings

To determine whether the HIVE and BREVE are degraded through 26S/UPS, coding sequences of *HIVE* and *BREVE* were cloned into pCAMBIA1300 containing firefly *luciferase* (LUC) reporter, to generate pLUC-HIVE or pLUC-BREVE. The pCAMBIA2301 containing the *GUS* reporter was used as internal control. The plasmids were separately mobilized into *A. tumefaciens* GV3101 and *C. roseus* seedlings were infiltrated using FAST method as previously described. After three days inoculation at NT, seedlings were transferred to cold room (4℃) for different time. Samples were collected for LUC and GUS activity assay conducted as previously described (Pattanaik et al., 2010).

#### Arabidopsis transformation, cold treatment, and freezing tolerance assays

The pCAMBIA1300 plasmids containing *HIVE* were used for Arabidopsis transformation by floral dip method (Clough & Bent, 1998). *HIVE* was transformed to homozygous *ice1-2* mutant plants (Kanaoka et al., 2008). T3 homozygous transgenic plants were used in this study. For cold treatment, three-week-old Arabidopsis seedlings were treated with 4°C for 1 hour. The whole seedlings were collected for RNA isolation. For freezing assays, three-week-old Arabidopsis seedlings were treated with -10°C for 2 hours and then kept at 4°C in the dark for 12 hours. The seedlings were then transferred to normal conditions for 4 days and survival rate was recorded.

#### Yeast one-hybrid and two-hybrid assays

To determine the interaction between the intermediate factor and *D4H* promoter, a Y1H assay was conducted. The full-length cDNA of *CrMYBR2* was cloned in pGADT7 and truncated *D4H* promoter was cloned in pHIS2. *pGADT7-CrMYBR2* and *pHIS2-proD4Hs* were transformed into yeast strain Y187. The co-transformed Y187 yeast strain was cultured on Trp/Leu/His medium supplemented with 50 mM and 100 mM 3-AT.

The full-length cDNAs of *HIVE* or *BREVE* were cloned in pAD-GAL4-2.1; full-length of cDNA of *CrSnRK2.6*, *CrSnRK2.7*, or bHLH domains of *HIVE* and *BREVE* were cloned in pBD-GAL4 Cam (Stratagene, USA). Different combinations of bait and prey plasmids were transformed into yeast strain AH109 using PEG/LiCl method (Clontech, USA), and the transformants were selected on synthetic dropout (SD) medium lacking leucine and tryptophan (-leu-trp). Transformed colonies were then dropped on SD medium lacking histidine, leucine, and tryptophan (-his-leu-trp) or adenine, histidine, leucine, and tryptophan (-ade-his-leu-trp) to check protein-protein interaction. Due to self-activation of the full-length TFs in yeast cells, TF fragments without activation domains but with bHLH domains were used in this assay.

#### RNA isolation, cDNA synthesis and quantitative RT-PCR

RNA isolation, cDNA synthesis and quantitative real-time PCR (RT-qPCR) were performed as previously described (Liu et al., 2019). Primers used in RT-qPCR and other primers used in this study are listed in Table S5.

#### Alkaloid extraction and analysis

Extraction and analysis of alkaloids from *C. roseus* samples were performed as described previously (Liu et al., 2019). The concentrations of the alkaloids were calculated using a standard curve.

#### Statistical analysis

In insect bioassay, statistical analysis was conducted in JMP^®^ PRO, version 14.0.0 (SAS Institute Inc., Cary, NC, US). Data normality was checked before analysis using the Shapiro-Wilk test (p > 0.05). For body weight, a two-way ANOVA was used with treatment and concentration and interaction between treatment and concentration as factors. One-way ANOVA was conducted in comparisons in effects of treatment and concentration for each time point. Tukey-Kramer HSD test was used to conduct comparisons for all pairs of treatments and control. Body weight in 48h were transformed by log(x) to meet the assumption of homogeneity of variance (Brown-Forsythe test, P > 0.05). Non-parametric analysis was used to analyze mortality data by Kruskal-Wallis test, followed by student’s t-test (extremely small sample size with large effect size (De Winter, 2013)) for all comparisons of each pair of treatments in mortality at each concentration due to the data was not and could not be transformed to meet the assumption of normal distribution. ANOVA was used to check the start area of all used leaves. Leaf consumed area was analyzed by Kruskal-Wallis test. The Wilcoxon signed-rank test was subsequently conducted to determine the differences among different treatments at different concentrations. In transcriptional experiments, statistical significance was calculated using Student’s *t* test (* P < 0.05 and ** P < 0.01) or ANOVA with Tukey’s honestly significant difference (HSD) test (P < 0.05). Error bars represented standard deviation (SD).

## Acknowledgments

We thank Megan Combs (Department of Civil Engineering and Environmental Research Training Laboratories, University of Kentucky) for assistance on LC-MS/MS analyses. This work is partially supported by the Harold R. Burton Endowed Professorship to L.Y., the C. W. Kearns, C. L. Metcalf, and W. P. Flint Endowed Chair Professorship in Insect Toxicology to X.Z., and National Key Crop Research Institute, Guangdong Academy of Agricultural Sciences/Guangdong Provincial Key Laboratory of Crop Genetic Improvement Open Research Fund (NYGC202202).

## Author Contributions

L.Y., Y.L., B.P., X.Z., YW and S.P. designed the research; Y.L., B.P., J.S., S.K.S., X.L., Y.L. and S.P. performed experiments; Y.L., J.S., S.K.S., X.Z., S.P. and L.Y. analyzed data; Y.L., S.P., X.Z., B.P. and L.Y. wrote the paper with inputs from all authors.

## Declaration of interests

The authors declare no conflict of interest.

## Lead Contact Website

https://ktrdc.ca.uky.edu/person/ling-yuan

https://ktrdc.ca.uky.edu/person/sitakanta-pattanaik

Xuguo (Joe) Zhou | School of Integrative Biology | UIUC (illinois.edu)

## Notes

### Competing Interest Statement

The authors have declared no competing interest.

