## Supplemental Figures and Tables for "Transcriptional reprogramming deploys a compartmentalized “timebomb” in *Catharanthus roseus* to fend off chewing herbivores"

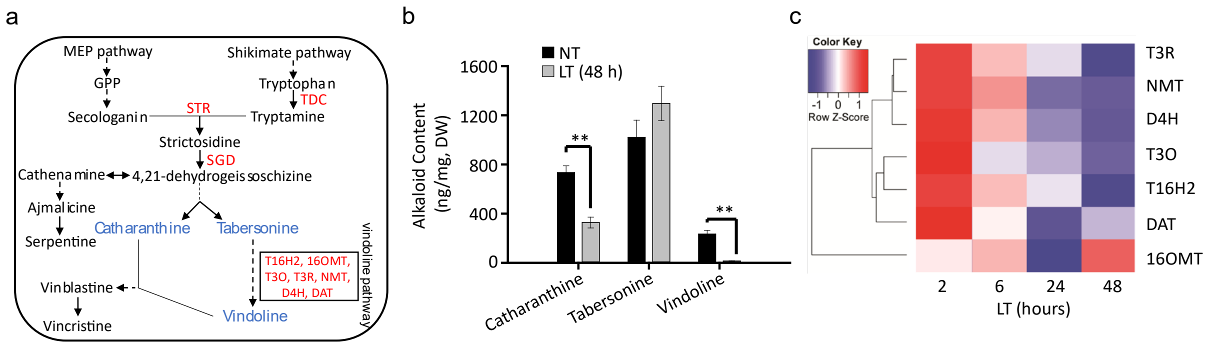


**Supplementary Figure 1: Vindoline biosynthesis is suppressed by low temperature in *C. roseus* seedlings.**

**a** A simplified diagram of the TIA biosynthetic pathway in *C. roseus*. 16OMT, 16-O-methyltranferase; D4H, desacetoxyvindoline-4-hydroxylase; DAT, deacetoxyvindoline-4-acetyletransferase; NMT, N-methyltransferase; SGD, strictosidine-β-D-glucosidase**;** STR, strictosidine synthase; T3O, tabersonine 3-oxygenase; T3R, tabersonine 3-reductase; T16H2, tabersonine 16-hydroxylase 2; TDC, tryptophan decarboxylase. Solid arrows indicate single enzymatic steps while dotted arrows indicate multiple enzymatic steps. **b** Concentrations of catharanthine, tabersonine, and vindoline in *C. roseus* seedlings. Normal temperature (NT) grown seedlings were either maintained at NT (control) or exposed to low temperature (LT; 4$^{o}$C) for 48 hours. Alkaloids were extracted and analyzed by LC-MS/MS, and the concentration of each alkaloid was estimated based on peak areas compared with standards. DW, dry weight. The values represent means ± SD from three biological replicates. Statistical significance was calculated using Student's *t* test (** *p* <0.01). **c** Relative expression of seven vindoline pathway genes (*T16H2*, *16OMT*, *T3O*, *T3R*, *NMT*, *D4H*, and *DAT*) in *C. roseus* seedlings exposed to LT for different lengths of time (2, 6, 24, 48 hours). Gene expression was measured by RT-qPCR and visualized using a heatmap generated with the R package.

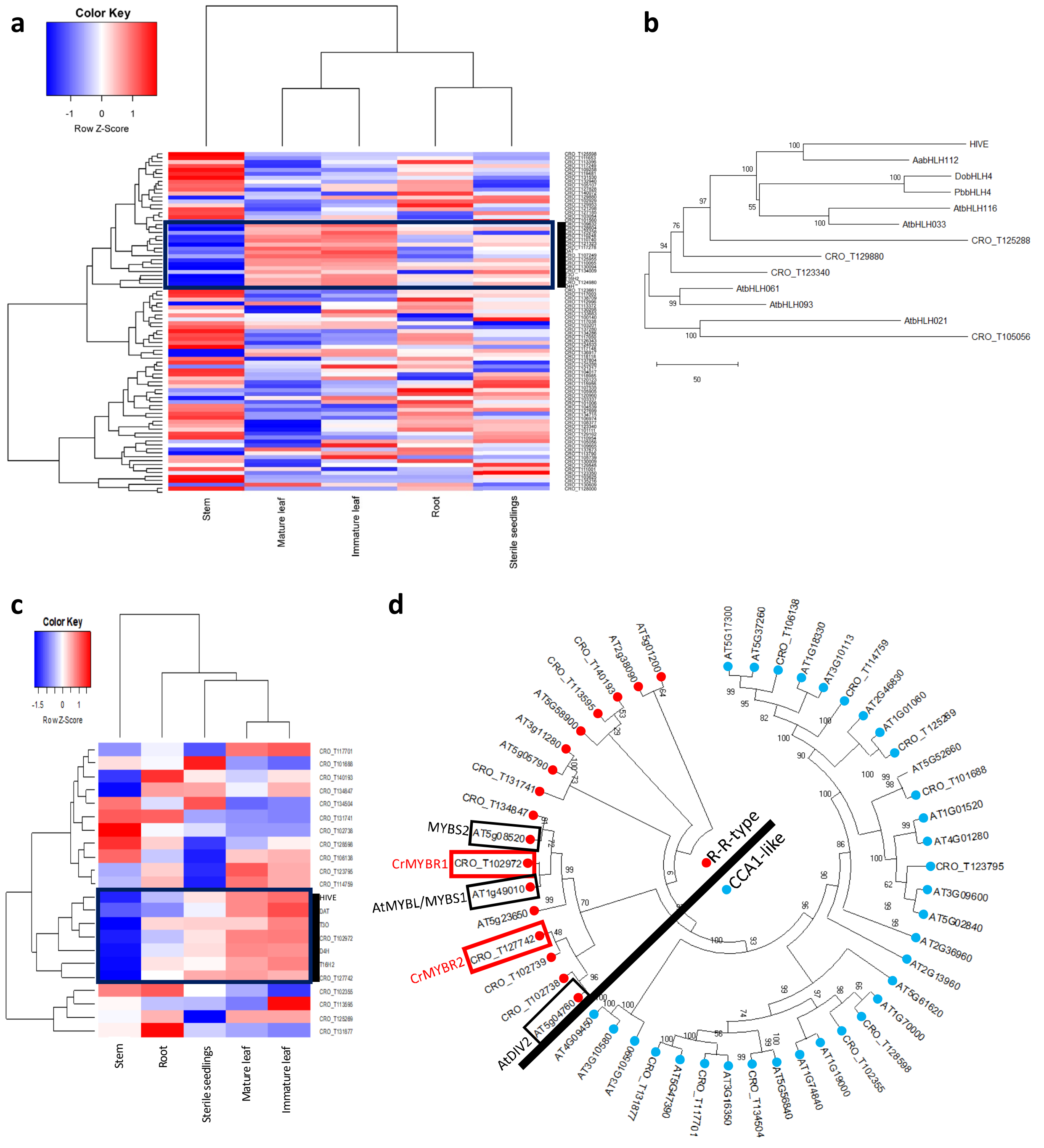


**Supplementary Figure 2: Identification of novel bHLH and MYBR regulators of vindoline biosynthesis.**

**a** Co-expression analysis of *CrbHLH* genes with vindoline pathway genes in five different *C. roseus* tissues. Hierarchical clustering and heat-map showing that 13 bHLH genes (Supplementary Table 1) are clustered with the four cold- and herbivory-regulated vindoline pathway genes. **b** Phylogenetic analysis of selected subgroup IIIb bHLH proteins from Arabidopsis, *C. roseus, Artemisia annua, Dendrobium officinale*, and *Phalaenopsis bellina*.

AtbHLH021 and CRO_105056 from subgroup IIIa were used as the outgroup. **c** Co-expression analysis of R-R-type and CCA1-like MYBs with *HIVE* and the vindoline pathway genes in five *C. roseus* tissues. Hierarchical clustering and the heat-map show that two R-R-type MYBs, CRO_T102971 and CRO_T127742, are clustered with *HIVE* and the four cold-regulated vindoline pathway genes. **d** Phylogenetic analysis of R-R-type and CCA1-like MYBs from *C. roseus* and Arabidopsis. The amino acid sequences of R-R-type and CCA1-like MYBs from *C. roseus* and Arabidopsis was used to construct a phylogenetic tree. The R-R-type MYBs are indicated by red dots while the CCA1-like MYBs are indicated by blue dots. *C. roseus* MYBR1 and MYBR2 are highlighted with red boxes and the previously reported Arabidopsis R-R-type factors are highlighted with black boxes. The phylogenetic trees were constructed using the Neighbor-Joining (N-J) method as implemented in MEGA5.1 software. The statistical reliability of individual nodes was assessed by bootstrap analysis with 1,000 replications. Scale bars represent substitutions per amino acid position.

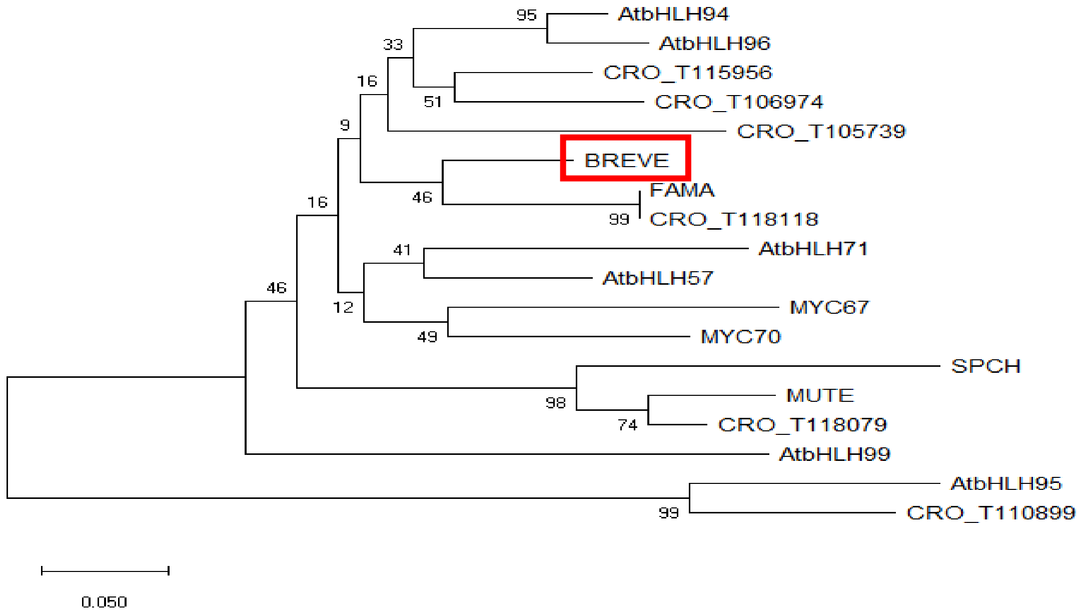


**Supplementary Figure 3: Phylogenetic analysis of BREVE.**

Phylogenetic analysis of the subgroup Ia bHLH TFs in Arabidopsis and *C. roseus*. The phylogeny was reconstructed and visualized using the Neighbor-Joining (N-J) method in MEGA5.1 software. The statistical reliability of individual nodes on the tree was assessed by bootstrap analysis with 1,000 replications. The scale bar represents substitutions per amino acid position. BREVE is highlighted with a red box. AtbHLH95 and CRO_110899 from subgroup Ib were used as the outgroup.

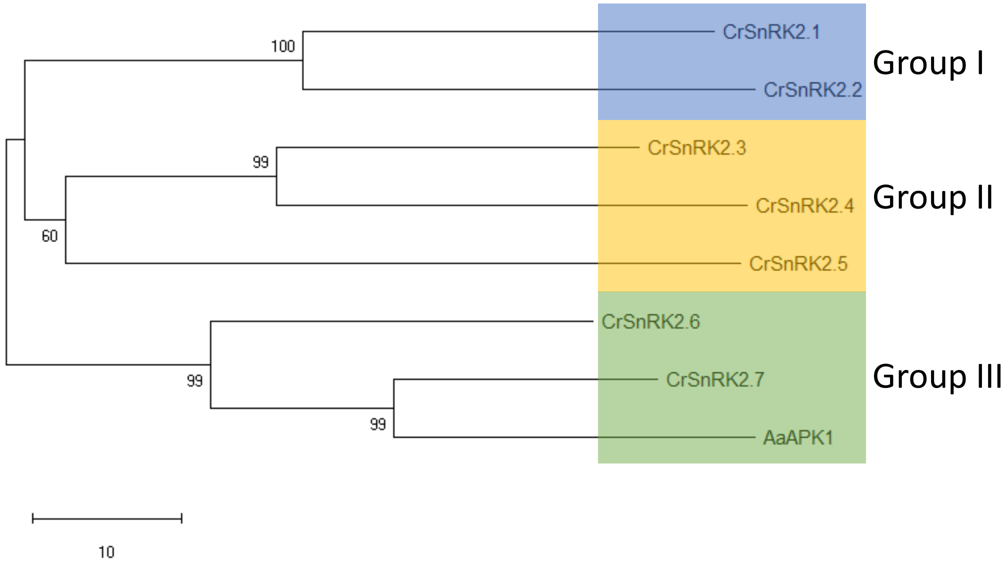


**Supplementary Figure 4: Phylogenetic analysis of *C. roseus* SnRK2 kinases.**

Phylogenetic analysis of SnRK2 subfamily kinases from *C. roseus* and AaAPK1. The phylogenetic tree was constructed from an alignment of the amino acid sequences of SnRK2s using the Neighbor-Joining (N-J) method in MEGA5.1 software. The statistical reliability of individual nodes was assessed by bootstrap analysis with 1,000 replications. The scale bar represents substitutions per amino acid position.

**Supplementary Table 1:** Thirteen *C. roseus* bHLHs co-expressed with vindoline pathway genes (See also Supplementary Fig. 2a). Genes are sorted by subgroup.

| Transcript ID | Subgroup |
|---|---|
| CRO_T124980 | Ia |
| CRO_T110248 | IIIb |
| CRO_T117278 | IIIc |
| CRO_T125258 | IIId |
| CRO_T125955 | IVc |
| CRO_T110065 | VIIb |
| CRO_T107249 | VIIb |
| CRO_T130094 | VIIb |
| CRO_T134009 | VIIIb |
| CRO_T110740 | XII |
| CRO_T121323 | XII |
| CRO_T128604 | XII |
| CRO_T109632 | XII |

**Supplementary Table 2:** The cis-elements in proD4Hs predicted by PlantRegMap.

| Motif | Family | Position | Strand | Matched sequence |
|---|---|---|---|---|
| cra_locus_5807 | C2H2 | 23-37 | - | TAATAAAGACAATTA |
| cra_locus_17076 | CPP | 284-296 | + | GAAAATTTTAAAC |
| cra_locus_21919 | CPP | 67-76 | + | AATTTGAATT |
| cra_locus_35394 | Dof | 26-46 | + | TTGTCTTTATTAACTCTCTTT |
| cra_locus_35394 | Dof | 239-259 | - | TTTATTTTTTTCCTTAATATT |
| cra_locus_3219 | MYB-related | 200-211 | - | AGATATTATATT |
| cra_locus_3605 | YABBY | 133-142 | - | CAATAATAAC |

**Supplementary Table 3:** The cis-elements in proCrMYBR2ss predicted by PlantRegMap.

| Motif | Family | Position | Strand | Matched sequence |
|---|---|---|---|---|
| cra_locus_5787 | bHLH | 217-227 | - | CGCAAGTTGCA |
| cra_locus_4348 | bHLH | 217-227 | + | TGCAACTTGCG |
| cra_locus_4179 | bHLH | 218-226 | + | GCAACTTGC |
| cra_locus_14386 | C3H | 400-413 | - | AGTCAAAAAGGCGA |
| cra_locus_86428 | Dof | 69-89 | - | GGAAAACACTAAAAGGACTAA |
| cra_locus_9747 | EIL | 129-149 | + | TTGTCTATATCCAATCCTTCT |
| cra_locus_5090 | HD-ZIP | 301-315 | - | AAAAATAATGATAAT |
| cra_locus_6373 | MYB | 128-148 | - | GAAGGATTGGATATAGACAAG |
| cra_locus_6581 | MYB | 32-50 | - | AGCATACAACTACATTCAC |
| cra_locus_6071 | Trihelix | 368-381 | - | AATTTTACCATGAT |
| cra_locus_17159 | WOX | 107-114 | - | GCAATCAA |
| cra_locus_3605 | YABBY | 301-310 | - | TAATGATAAT |

**Supplementary Table 4:** Design of the TIA feeding assay.

| Concentration (μmol/ml) | chemical | No. of replicates |
|---|---|---|
| 2.19 | vindoline | 10 |
| 0.219 | vindoline | 10 |
| 0.0219 | vindoline | 10 |
| 2.19 | catharanthine | 10 |
| 0.219 | catharanthine | 10 |
| 0.0219 | catharanthine | 10 |
| 2.19 | vindoline + catharanthine | 10 |
| 0.219 | vindoline + catharanthine | 10 |
| 0.0219 | vindoline + catharanthine | 10 |
| control | methanol | 10 |

**Supplementary Table 5:** Primers used in this study.

| primer | sequence (5'-3') |
|---|---|
| | **For RT-qPCR** |
| PRS9-qPCR-F | GAGGGCCAAAACAAACTTGA |
| RPS9-qPCR-R | CCCTTATGTGCCTTTGCCTA |
| T16H2-qPCR-F | GATCAACTCACAGTGGCAGTC |
| T16H2-qPCR-R | GACTTGAGGACTTGTGATTGGC |
| 16OMT-qPCR-F | GTGGATTCTCCATGACTGGAACGA |
| 16OMT-qPCR-R | GATTATCACCTTTCCACCCTTCGC |
| T3O-qPCR-F | TTTGCCATTTGGTGCCGGAAGA |
| T3O-qPCR-R | CTGGGAGTTGCCAGTTGAAATGGT |
| T3R-qPCR-F | CGCGAGTACGGGTGGAAGTATAAA |
| T3R-qPCR-R | CGGGGATAACCTCAACATCTGCAA |
| NMT-qPCR-F | TTTGGCTTCATTATTGATGTTACC |
| NMT-qPCR-R | CTGGTGTCTTTAATAGGTTGTTCG |
| D4H-qPCR-F | GGCTGGTGTAAAAGGGATTGT |
| D4H-qPCR-R | TCTCACGCCGTATTTCTGAAT |
| DAT-qPCR-F | ATCGGTTGAGACAGAGACACTCTC |
| DAT-qPCR-R | GATACGCACGTTTGGTATATGTTTT |
| IPRT1-qPCR-F | TGCCATTATCAAGAACGTTGAG |
| IPRT1-qPCR-R | TGTTATCCTCGTCCAATTCCTC |
| HIVE-qPCR-F | TGGGAAAGATCAAGAAACTGGT |
| HIVE-qPCR-R | CATCCTTGTTCTCCAAAACTCC |
| CrMYBR1-qPCR-F | TTCAGGTTTAGGCCATGATTCT |
| CrMYBR1-qPCR-R | TCCCCTTTTCCAAACTTGTCTA |
| CrMYBR2-qPCR-F | TGTACCAGCAGAAAGAGGAACA |
| CrMYBR2-qPCR-R | TCGTCCAGAGTATGCCCTTATT |
| BREVE-qPCR-F | ATTCAGCGGATGACATAGCG |
| BREVE-qPCR-R | GCAGCAGCTTGACGATGAATC |
| CrSnRK2.7-qPCR-F | ATGATGTGCAACCAGTTTCAAG |
| CrSnRK2.7-qPCR-R | CATCATCCATTAGGTCCAGGTT |
| | **For transient gene overexpression** |
| HIVE-BamHI-F | CGCGGATCCATGTTATCGAGAGTTAACAGCATGG |
| HIVE-XbaI-R | ACGTTCTAGATTAAACCATCCCTTGGAAGC |
| CrMYBR2-BamHI-F | CAGTGGATCCATGGAAACTGATCGGACATG |
| CrMYBR2-XbaI-R | CAGTTCTAGATTACAGAGGCAATGGATATGC |
| BREVE-BamHI-F | CGCGGATCCCATGGCTTTAGAAGCCCTTTCTTCC |
| BREVE-XbaI-R | ACGTTCTAGATAACAAGTACGGGGTGGCGGGGGT |
| CrSnRK2.7-BamHI-F | ACTGGGATCCATGGATCGGGCGCCGATAAC |

| | |
|---|---|
| CrSnRK2.7-XbaI-R | ACTGTCTAGATTACATTGCATATATGACTTCTCC |
| | **For VIGS constructs** |
| CrChlH-vigs-F | ACTGGGTACCAGTTGCCACACTAGTTAATATTGCTGC |
| CrChlH-vigs-R | ACTGCTCGAGGCATGGATATTCTTTCCCGTTGGC |
| HIVE-vigs-F | ATGGGTACCTGCTTGAAGTTGAAGATGAGGA |
| HIVE-vigs-R | ATGCTCGAGAATCACTGAATCCGCCATTAAC |
| BREVE-vigs-F | ATGGGTACCTCAAATTCTGCTGCAACTTTGG |
| BREVE-vigs-R | ATGCTCGAGAATTTAGTTGACTTGCACAAA |
| | **For promoter isolation and cloning** |
| proT16H2-EcoRI-F | ACTGGAATTCACCCTTTTATAGGAGGAAAGC |
| proT16H2-XhoI-R | ACTGCTCGAGGAGGTAGAGAAAGTTGACC |
| proT3O-EcoRI-F | ACTGGAATTCCATGCCAAATGCCAAAGTATAGG |
| proT3O-XhoI-R | ACTGCTCGAGAATGAATATTTTTTGTTGGTTCGCTA |
| proD4H-EcoRI-F | ATGGAATTCCCAGGTTGGGTGAAGGTGC |
| proD4H-XhoI-R | ACTGCTCGAGTTTTCTTTCTTGCTCAGAATTTGGAAG |
| proDAT-EcoRI-F | ATGGAATTCCATTATAGGTGTATCTTCCCAC |
| proDAT-XhoI_R | ACTGCTCGAGTTTGCTTGCTGTTATATATTCAGACC |
| proD4Hs-EcoRI-F | GGGAATTCCGAGTTATCCTTATTAATTTC |
| proD4Hs-XhoI-R | GGCTCGAGTGTTTAAAATTTTCTTCTATA |
| proCrMYBR2-EcoRI-F | ACTGGAATTCGAATTATGATGTGGCAAGTGATTG |
| proCrMYBR2s-EcoRI-F | ACTGGAATTCTCACAACCAAGGATGTTTG |
| proCrMYBR2ss-EcoRI-F | ACTGGAATTCTGGAAAATCCTCGTTTGC |
| proCrMYBR2-SalI-R | ACTGGTCGACGTCCGATCAGTTTCCATTCG |
| | **For site-directed mutation** |
| proD4Hm1-F | GAAAATTTTCCCCCCAAAATAAAATTTTTTTAATTTTTTCGAGTTA TCCTT |
| proD4Hm1-R | TTATTTTGGGGGGAAAATTTTCTCTCTCTAAAACAATTACC |
| proD4Hm2-F | CTTGGCAAAAAAATGATATCATCTCATAGTTTCTGTTTTC |
| proD4Hm2-R | TGATATCATTTTTTGCCAAGACTTAGGATTTTTATAGG |
| proD4Hm3-F | AATATAAAAAAATTTTATTATAAAAATTTAATAAAAATAAATATTA AGG |
| proD4Hm3-R | TAAAATTTTTTATATTAATATGTTACATAGATGGTTAC |
| proCrMYBR2m-F | TGTGCCCCCCGCGAAATTGGTTACAAGTTTATATTG |
| proCrMYBR2m-R | TTCGCGGGGGGCACAACAAAGGCTGAA |
| proCrCBF1mE-F | TGACTCCATCAAAAACAACTAGCCATTCCTTTAAATATT |
| proCrCBF1mE-R | GAATGGCTAGTTGTTTTTGATGGAGTCAAATCAAAGTGAGACAA A |
| proCrCBF1mEL-F | AACTGCCCCCCGCCATTCCTTTAAATATTAAAATAAAAAAATACG |
| proCrCBF1mEL-R | AATGGCGGGGGGCAGTTGATGGAGTCAAATCAAAG |
| proCrCBF1MUT-F | CCATTCCGTGTGAAAATCCCGATCGTATCCTTCCTTTCTTCTC |
| proCrCBF1MUT-R | TACGATCGGGATTTTCACACGGAATGGGGATATGATAAGAAGC |

| | |
|---|---|
| $HIVE^{S348A}$-F | TATGATGCAGATGACTTAACAGAGAATACAAAG |
| $HIVE^{S348A}$ -R | TCATCTGCATCATAGTTCAGATTAGACCCATC |
| $HIVE^{S348D}$ -F | TATGATGACGATGACTTAACAGAGAATACAAAG |
| $HIVE^{S348D}$ -R | TCATCGTCATCATAGTTCAGATTAGACCCATC |
| | **For yeast-one hybrid** |
| proD4Hs-EcoRI-F | GGGAATTCCGAGTTATCCTTATTAATTTC |
| proD4Hs-SacI-R | GGGAGCTCTGTTTAAAATTTTCTTCTATA |
| CrMYBR2-BamHI-F | CAGTGGATCCATGGAAACTGATCGGACATG |
| CrMYBR2-XbaI-R | CAGTTCTAGATTACAGAGGCAATGGATATGC |
| | **For yeast-two hybrid** |
| HIVE-BamHI-F | CGCGGATCCATGTTATCGAGAGTTAACAGCATGG |
| HIVE-XbaI-R | ACGTTCTAGATTAAACCATCCCTTGGAAGC |
| BREVE-BamHI-F | CGCGGATCCCATGGCTTTAGAAGCCCTTTCTTCC |
| BREVE-XbaI-R | ACGTTCTAGATAACAAGTACGGGGTGGCGGGGGT |
| HIVE(bHLH)-EcoRI-F | CGCGAATTCAGCACAGTCACCGGAGAGG |
| HIVE(bHLH)-SalI-R | TGCGTCGACCTATCTTCCTTCACGTAACCTGACT |
| BREVE(bHLH)-EcoRI-F | CGCGAATTCAGGAGACAGAGAGTATGTAAG |
| BREVE(bHLH)-SalI-R | TGCGTCGACTCAACATGTGTATTGAGGGTATGTG |
| CrSnRK2.6-MfeI-F | ACTGCAATTGATGGATCGAGCGGCTGTGAC |
| CrSnRK2.6-XhoI-R | ACTGCTCGAGTTACATCGCGTAGATTATCTCTCCGCTG |
| CrSnRK2.7-EcoRI-F | ACTGGAATTCATGGATCGGGCGCCGATAAC |
| CrSnRK2.7-XhoI-R | ACTGCTCGAGTTACATTGCATATATGACTTCTCC |
